# Dysregulation of PAX5 causes uncommitted B cell development and tumorigenesis in mice

**DOI:** 10.1101/2021.01.29.428877

**Authors:** Brigette Boast, Kaiyue Helian, T. Daniel Andrews, Xi Li, Vicky Cho, Adria Closa, Henry J. Sutton, Joanne H. Reed, Hannes Bergmann, Carla M. Roots, Mehmet Yabas, Nadine Barthel, Sofia A. Omari, Clara Young, Lisa A. Miosge, Eduardo Eyras, Stephen L. Nutt, Nadine Hein, Katherine M. Hannan, Ian A. Cockburn, Christopher C. Goodnow, Anselm Enders

## Abstract

PAX5 is the master transcription factor controlling B cell identity. In humans, mutations in *PAX5* account for 30% of B cell acute lymphoblastic leukemia (B-ALL) cases. Investigating the causal effects of *PAX5* mutations has however been difficult due to the premature lethality of *Pax5*^−/−^ mice. Here we describe a novel mouse strain with a premature STOP mutation in *Pax5* (Y351*) that produces a truncated protein and reduction in protein function, yet still allows for some B cell development to occur. A population of uncommitted and multipotent CD19^+^B220^−^ B cells develops in the bone marrow of homozygous mice leading to the development of B-ALL. We show that the tumors frequently acquire secondary mutations in *Jak3*, and *Ptpn11* highlighting key pathways interacting with PAX5 during malignant transformation. Analysis of the PAX5^Y351*^ mice provide insight not only into the functional consequence of reduced PAX5 activity on B cell development and identity, but also provides an avenue in which to study PAX5-driven B-ALL in mice.

**One Sentence Summary:** Reduction in PAX5 function in mice induces the development of uncommitted B cells that have multipotent and malignant potential.

## Introduction

Paired box gene 5 (PAX5) is often referred to as the guardian of B cell identity due to its crucial role in establishment and maintenance of the B cell transcriptional program (*1*). PAX5 establishes B cell identity early in lymphoid development by acting in a regulatory network with E2A, EBF1, and IKZF1 to push uncommitted lymphoid progenitors into a committed pro-B cell in the bone marrow (*2–4*). It then initiates transcription of B cell-specific genes and repression of non-B cell genes (*4–8*) and facilitates *Igh* V_H_-DJ_H_ recombination in pro-B cells, (*9–11*) allowing for progression through to pre-B cells. PAX5 expression is then maintained throughout the rest of development until it is lost during terminal differentiation into plasma blasts and plasma cells (*12, 13*).

PAX5-deficient mice (*Pax5*^−/−^) have a complete absence of mature B cells in the periphery due to a block of development at the pro-B cell stage in the bone marrow (*11, 14*). In contrast to wildtype pro-B cells, PAX5-deficient pro-B cells have a broad multi-lineage potential with the ability to dedifferentiate into all hematopoietic cell types, with the exclusion of B cells, both *in vitro* (*15*) and *in vivo* (*16–19*). This shows that PAX5 is crucial for the commitment to the B cell lineage in pro-B cells (*15*).

*Pax5*^−/−^ mice fail to thrive and die shortly after weaning, possibly due to a dependence of PAX5 on the development of the posterior midbrain (*14*). This complicates any analyses into the effects of PAX5-deficiency on B cell development in adult mice. To combat this issue, conditional deletions of PAX5 using *Cd19-Cre* and *Mx1-Cre* (*10*) have been developed to circumvent the need for PAX5 in the nervous system, but have been inefficient at exclusively deleting *Pax5* in 100% of B cells. To further this work, the same group developed a system where *Pax5* was deleted in mature B cells *in vitro* that were then transferred into sub-lethally irradiated *Rag2*^−/−^ mice (*20*). This showed that deleting *Pax5* in mature B cells reverts their commitment from the B cell lineage, causing dedifferentiation back to pluripotent progenitors and subsequent development of T cells and myeloid cells from the same previously committed B cells (*20*). This suggests that PAX5 is crucial not only for driving B lineage commitment in pro-B cells, but also to maintain lineage commitment throughout the life of the B cell.

In addition to its role as the major B cell transcription factor, PAX5 is also an essential tumor suppressor in both humans and mice. It is the most commonly mutated gene in B cell acute lymphoblastic leukemia (B-ALL) with approximately 30-40% of all B-ALL cases carrying a mutation in *PAX5* (*21, 22*). Extensive work has been done to identify different *PAX5* mutations in human B-ALL, with many of these mutations resulting in either a loss or reduction of PAX5 function (*21, 23, 24*). Thus far, there has only been one report of germline heterozygous mutations in humans, resulting in familial pre-B ALL development with incomplete inheritance (*25*). This particular mutation (G183S) occurs in the octapeptide domain and results in a reduction in the transactivating capacity of PAX5 (*25*). In mice, multiple studies have demonstrated that hypomorphic *Pax5* mutations are potent drivers of B-ALL (*20, 26–28*). While mice lacking one or both copies of *Pax5* do not spontaneously develop B-ALL, when crossed to a constitutively active form of STAT5, haploinsufficiency of *Pax5* results in a faster onset of B-ALL compared with mice that only have the gain-of-function in STAT5 (*27*). Interestingly, when *Pax5* is conditionally deleted in B cells, aged mice will spontaneously develop a B cell-derived lymphoma (*20*). Additional work has been done to show that some of the mutations identified in *PAX5*-driven B-ALL in human cohorts, also induces the spontaneous development of CD19^−^B220^+^ B-ALL in heterozygous mice (*23, 29*).

Here we describe a novel mouse strain with a single point mutation in *Pax5* that leads to a premature stop codon at amino acid 351 at the C terminus. This Y351* mutation results in the stable expression of a truncated, hypomorphic version of PAX5, with homozygous mice surviving well into adulthood. Aged *Pax5*^*Y351*/Y351\**^ spontaneously develop precursor B-ALL with 100% penetrance. The *Pax5*^*Y351*/Y351\**^ mice provide an avenue to study the effects of reduced PAX5 function in 100% of B cells in adult mice and provide an important tool for studying the oncogenic events that occur prior to malignant transformation.

## Results

### Subhead 1: A Novel Pax5 Mutation

An *N-*ethyl-*N-*nitrosourea (ENU) mutagenesis screen identified several related mice with a very low proportion of B cells in their blood (Fig. 1, A and B). Analysis of exome sequencing data of founder G1 mice revealed an A to T substitution converting tyrosine 351 to STOP at the C terminal domain of PAX5. The premature stop codon is predicted to remove the entire 33 amino acids of the inhibition domain as well as the neighboring 8 amino acids of the transactivation domain (Fig. 1C). In contrast to *Pax5*^−/−^ mice, *Pax5*^*Y351*/Y351\**^ mice maintained low frequencies (1.5%) of circulating B cells, albeit severely reduced compared to wildtype (60%) (Fig. 1, A and B), indicating that the *Pax5*^*Y351*/Y351\**^ mutation does not behave as a null allele. A western blot of PAX5 expression on isolated CD19^+^ B cells from the bone marrow of *Pax5*^*wt/wt*^, *Pax5*^*wt/Y351*\*^, and *Pax5*^*Y351*/Y351\**^ mice confirmed that the Y351* mutation prematurely arrests PAX5 translation (Fig. 1D). *Pax5*^*Y351*/Y351\**^ B cells expressed a truncated PAX5 protein and *Pax5*^*wt/Y351\**^ B cells expressed both full-length and truncated PAX5 proteins (Fig. 1D). The Y351* mutation therefore does not lead to the absence of PAX5 protein and results in the expression of a truncated PAX5 protein. The truncated PAX5 protein is stably expressed, with peripheral B cells showing upregulation of PAX5 expression in both heterozygous and homozygous mice (Fig. 1E).

**Fig. 1.**
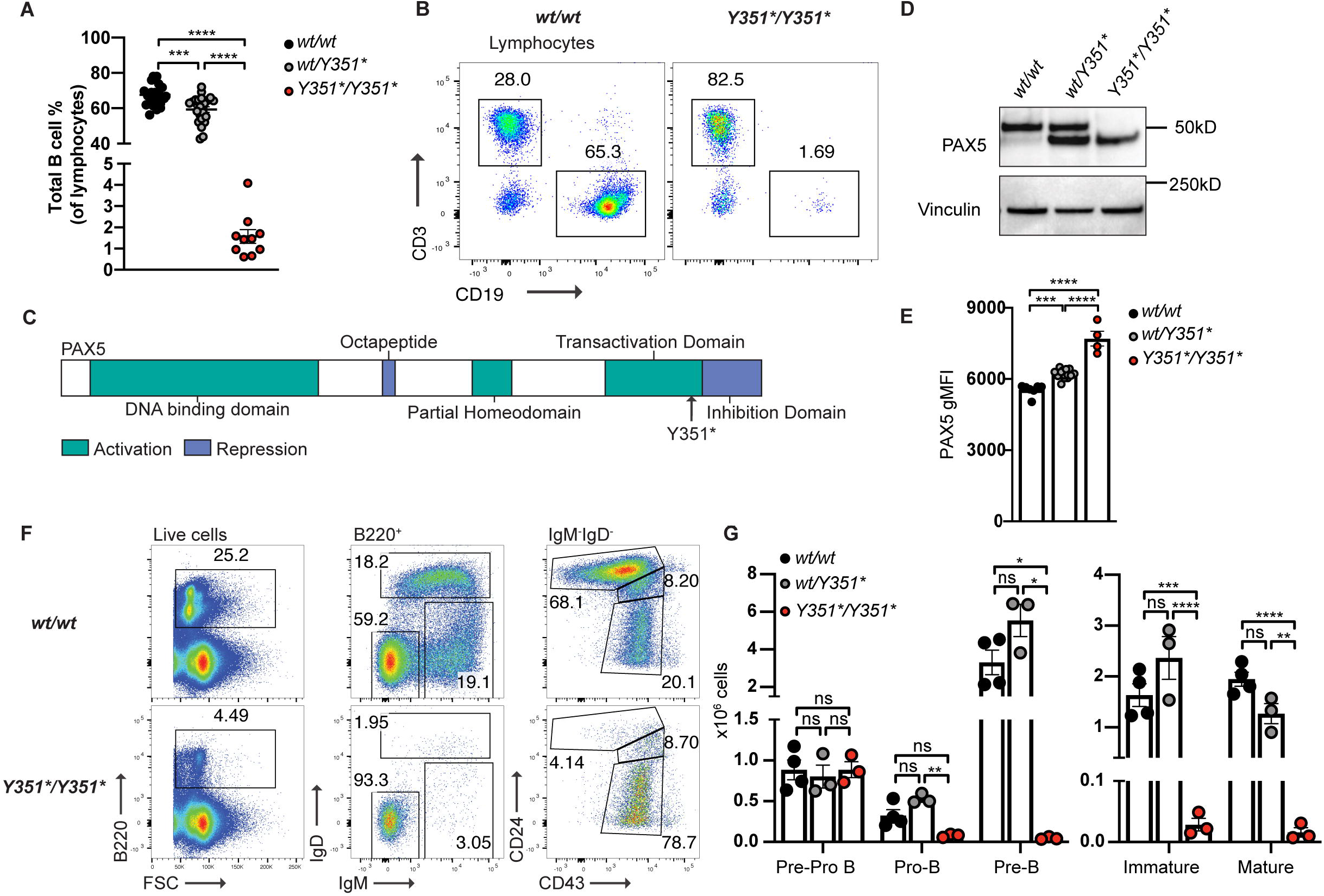
A novel truncating mutation in *Pax5* causes a severe reduction in the frequency of circulating B cells in homozygous mice. **(A)** Frequency of CD19^+^ B cells in the peripheral blood determined by flow cytometry. Data from one experiment and representative of ≥ five independent experiments. **(B)** Representative FACS plot of live lymphocytes from the blood showing B and T cell frequencies. **(C)** Schematic of PAX5 showing the location of Y351* mutation in relation to PAX5 functional domains. **(D)** Western blot of PAX5 expression from MACS-isolated CD19^+^ B cells from the bone marrow, representative of two independent experiments. **(E)** gMFI of PAX5 expression on B cells in the blood determined by flow cytometry, representative of three independent experiments. **(F)** Representative FACS plots of bone marrow gating strategy in 14-week-old littermates. **(G)** Total number (x10^6^) of B cell subsets in the bone marrow determined by flow cytometry, representative of three independent experiments. **(A, E, G)** Lines/bars show average per group ± SEM with individual mice indicated by circles. Asterisks indicate significance: ns P≥0.05, *P<0.05, **P<0.01,***P<0.001, ****P<0.0001 by one-way ANOVA with Tukey correction for multiple comparisons.

In contrast to *Pax5*^−/−^ mice, *Pax5*^*Y351*/Y351\**^ mice survive well into adulthood with a slight but significant reduction in weight compared to wildtype littermates (Fig. S1). Heterozygous mice maintained normal weight comparable to wildtype littermates (Fig. S1). This indicates that the *Pax5*^*Y351*/Y351\**^ mutation is not as severe as a complete null to cause premature death but does have some effects on the overall weight of homozygous mice. Overall, the presence of low frequencies of circulating B cells, the expression of a stable but truncated form of PAX5, and the effect on growth in *Pax5*^*Y351*/Y351\**^ mice indicates that although mutated, the expression of truncated PAX5 must retain some functional capacity that is sufficient for some B cell development to occur.

### Subhead 2: Reduced capacity for normal B cell development in Pax5*Y351\** mutants

To observe the effects of the Y351* mutation on B lymphopoiesis, we analyzed B cell development in the bone marrow and found a partial block at the pro-B cell stage in homozygous mice (Fig. 1, F and G). In agreement with the presence of low numbers of B cells in peripheral blood (Fig. 1A), this block was incomplete as we could identify low numbers of immature (IgM^+^IgD^−^) and mature (IgM^+^IgD^+^) B cells (Fig. 1G), suggesting that expression of a truncated form of PAX5 is sufficient for some cells to be able to successfully rearrange their BCR and complete maturation in the absence of full-length PAX5. Although heterozygous mice had a modest reduction of circulating B cells (Fig. 1A), we did not observe a significant block in early B cell development in the bone marrow (Fig. 1G).

To determine how terminal B cell maturation was affected by the *Pax5*^*Y351\**^ mutation, we analyzed splenic B cell populations by flow cytometry and found that total numbers of B220^+^ B cells were significantly decreased in homozygous and to a lesser extent in heterozygous mice (Fig. 2A). There was a significant reduction in follicular B (FoB) (CD23^+^CD21/35^low^) cells in both heterozygous and homozygous mice, with homozygotes having a much more severe reduction than heterozygotes (Fig. 2, B and C). Although *Pax5*^*Y351*/Y351\**^ B cells co-expressed IgM and IgD, the expression of IgD was decreased compared to *Pax5*^*wt/wt*^ FoB cells (Fig. 1D and Fig. S2). Heterozygotes had normal numbers of marginal zone B cells (MZB) (CD23^−^ CD21/35^high^) whilst homozygotes had significantly reduced numbers despite their increased frequency compared to FoBs (Fig. 2C and D). Despite a differential skew towards MZB frequencies, *Pax5*^*Y351*/Y351\**^ MZB cells appeared similar to wildtype cells with regards to CD1d and CD9 expression (Fig. 2E). Similar to the spleen, the lymph nodes in *Pax5*^*Y351*/Y351\**^ mice showed significantly reduced numbers of B cells (Fig. 2F). Mutant B cells in the lymph node appeared to be phenotypically similar to wildtype FoB cells with a slightly reduced CD23 and increased CD21 expression (Fig. 2G). IgD expression was also decreased on the surface of *Pax5*^*Y351*/Y351\**^ B cells in the lymph nodes (Fig. 2G). A similar observation was made when *Pax5* is deleted from mature B cells in the periphery (*10*) suggesting a role for PAX5 in the maintenance of IgD expression. Analysis of the peritoneal cavity revealed a preferential skew towards B1, and specifically B1a cells, in both heterozygous and homozygous mice (Fig. S3, A and B). This could be a result of differences in PAX5 dependency in the fetal liver compared to the adult bone marrow (*11*), suggesting that PAX5^Y351*^ promotes the development of self-renewing B1 cells prior to hematopoiesis in the bone marrow.

**Fig. 2.**
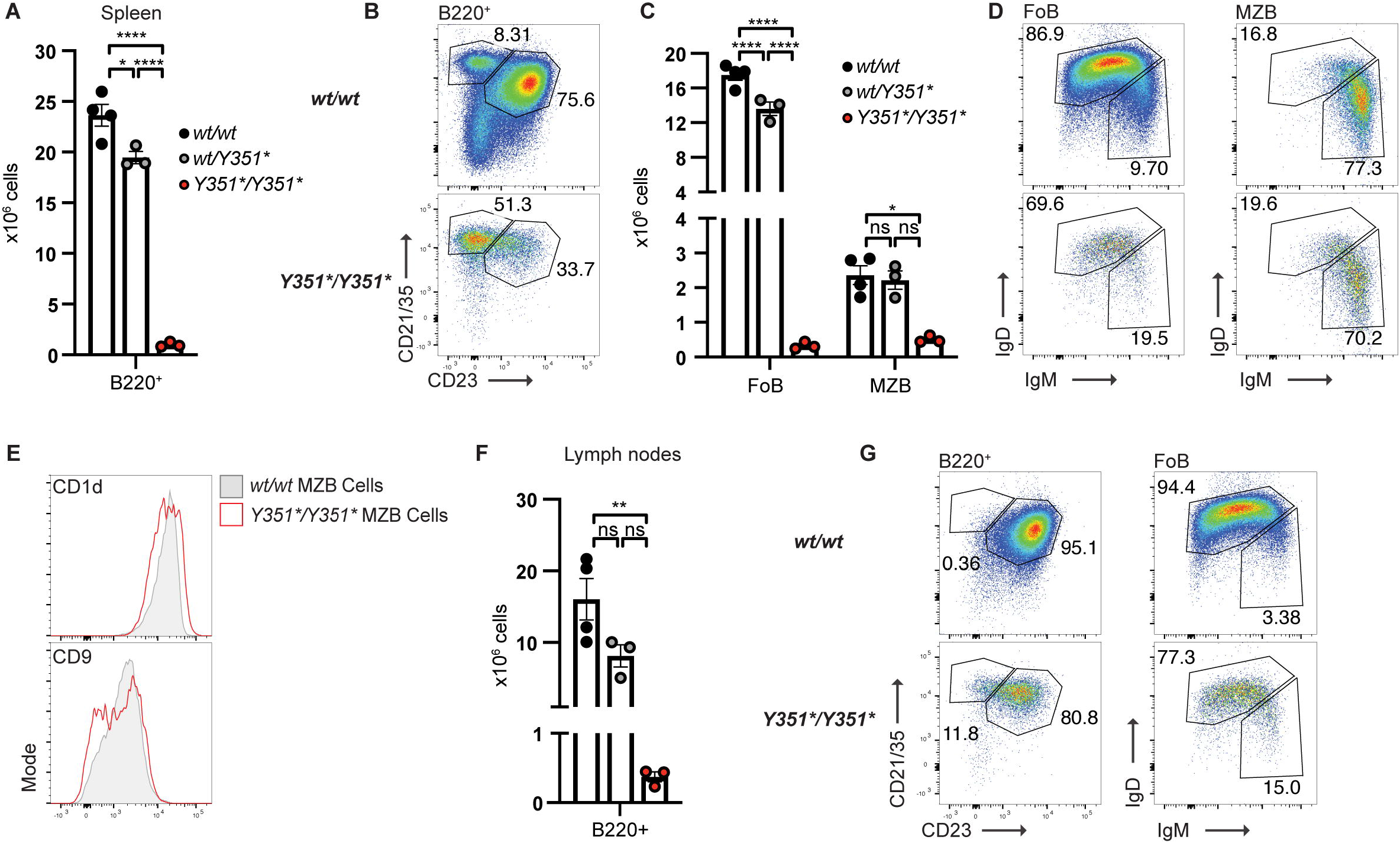
PAX5^Y351*^ causes a reduction in peripheral B cells in secondary lymphatic organs. **(A)** Total number (x10^6^) of B220^+^ B cells in the spleen. **(B)** Representative FACS plots of spleen gating strategy showing wildtype and homozygous mice, follicular B cells (FoB) gated CD23^+^CD21/35^low^, marginal zone B cells (MZB) gated CD23^−^CD21/35^hi^, pre-gated on live B220^+^ single cells. **(C)** Total number of FoB and MZB cells (x10^6^) in the spleen. **(D)** Representative FACS plots of IgM and IgD expression on B220^+^ FoB and MZB cells in the spleen of wildtype and homozygous mice. **(E)** Representative expression (gMFI) of CD1d and CD9 on wildtype and homozygous MZB cells. **(F)** Total number (x10^6^) of B220^+^ B cells in the lymph nodes. **(G)** Representative FACS plots of lymph node (2x inguinal and 2x brachial) gating strategy showing wildtype and homozygous mice, follicular B cells (FoB) gated CD23^+^CD21/35^low^ on live B220^+^ single cells. **(A-G)** Representative of three independent experiments. **(A, C, E)** Bars show average per group ± SEM with individual mice indicated by circles. Asterisks indicate significance: ns P≥0.05, *P<0.05, **P<0.01, ***P<0.001, ****P<0.0001 by one-way ANOVA with Tukey correction for multiple comparisons.

### Subhead 3: An alternate pathway of B cell development

In addition to the attenuation of normal B cell development in the bone marrow, we also observed an extra population of CD19^+^B220^−^ cells in the bone marrow, spleen, and lymph nodes of *Pax5*^*Y351*/Y351\**^ mice (Fig. 3A). Despite an overall reduction in total splenocyte number, the population of CD19^+^B220^−^ cells were greatly expanded in all tissues analyzed (Fig. 3B). Except for the absence of B220 expression, these CD19^+^B220^−^ cells in the bone marrow appeared to be phenotypically similar to pro-B cells (predominantly IgM^−^IgD^−^CD43^+^CD24^+^) (Fig. 3C), albeit with slightly increased CD43 and CD93 expression (Fig. 3, D and E). By contrast, in the spleen and lymph nodes, CD19^+^B220^−^ cells were negative for CD93 (Fig 3D) and positive for IgM and could be divided into IgM^hi^CD5^−^ and IgM^low^CD5^+^ (Fig. 3F). In wildtype mice, these cells represent B1b and B1a cells respectively. Furthermore, CD19^+^B220^−^ cells in the bone marrow and spleen maintained elevated expression of PAX5 suggesting an initial commitment to the B cell lineage (Fig. S4A).

**Fig. 3.**
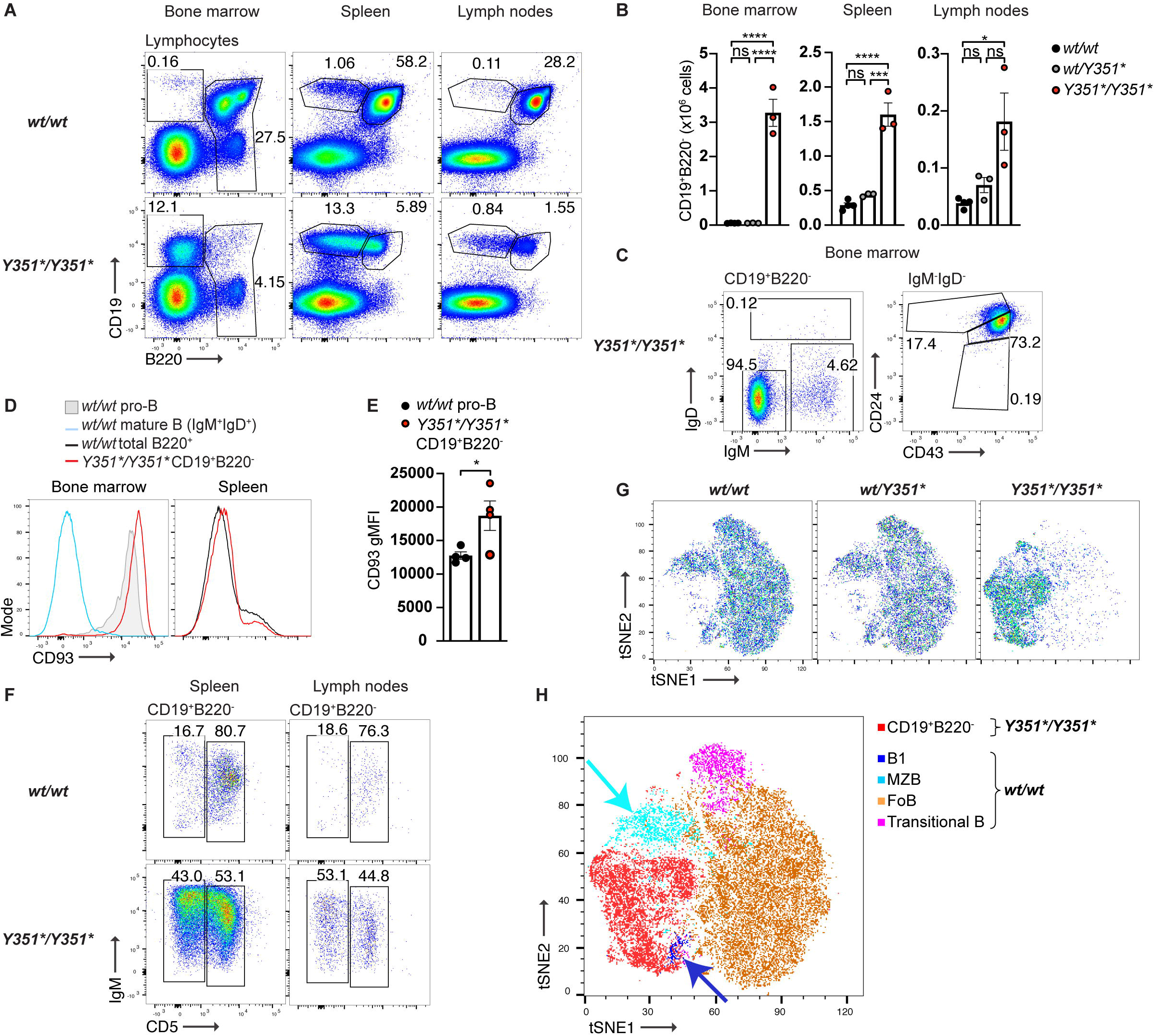
Emergence of an abnormal population of CD19^+^B220^−^ B cells in *Pax5*^*Y351*/Y351\**^ mice. Representative FACS plots showing the presence of CD19^+^B220^−^ cells in the bone marrow, spleen, and lymph nodes of homozygous mice that are not present in wildtype. Total cell numbers (x10^6^) of CD19^+^B220^−^ cells in the bone marrow, spleen, and lymph nodes. **(C)** Representative FACS plots of CD19^+^B220^−^ phenotype in the bone marrow, demonstrating likeness to wildtype pro-B cells using the gates set for wildtype precursor B cells similar to Figure 1F. **(D)** Representative histograms of CD93 expression in the bone marrow and spleen on *Pax5*^*wt/wt*^ mature (IgM^+^IgD^+^) B cells (blue line), *Pax5*^*wt/wt*^ pro-B cells (grey shaded), *Pax5*^*Y351*/Y351\**^ CD19^+^B220^−^ (red line), and *Pax5*^*wt/wt*^ total B cells (blue shaded), determined by flow cytometry. **(E)** Quantitative expression (gMFI) of CD93 on *Pax5*^*wt/wt*^ pro-B cells and *Pax5*^*Y351*/Y351\**^ CD19^+^B220^−^ cells from the bone marrow, determined by flow cytometry. **(F)** Representative FACS plots of the bone spleen and lymph nodes showing the IgM and CD5 expression on CD19^+^B220^−^ cells compared to wildtype B1 cells (CD19^+^B220^−^). **(G)** tSNE plots showing the overall likeness of B cells in the spleen in wildtype, heterozygous, and homozygous mice. Live B cells (CD19^+^) from each sample were down-sampled to 5000 cells and concatenated into one file. A tSNE analysis was run on the concatenated file using a panel of 11 different markers: CD9, CD23, CD19, CD21/35, CD1d, IgM, IgD, CD43, B220, CD5, CD11b. Each panel shows the result of the tSNE analysis per genotype; *wt/wt* n=4, *wt/Y351\** n=3, *Y351*/Y351\** n=3. **(H)** tSNE plot showing the clustering of similar B cell types based on their two-dimensional location. MZB: CD19^+^B220^+^CD21/35^hi^CD23^−^, FoB: CD19^+^B220^+^CD21/35^low^CD23^+^, B1: CD19^hi^B220^−^, Transitional B: CD19^+^B220^+^CD21/35^−^CD23^low^. *wt/wt* n=4, *Y351*/Y351\** n=3. **(B, E, F)** Bars show average per group ± SEM with individual mice indicated by circles. Asterisks indicate significance: ns P≥0.05, *P<0.05, **P<0.01, ***P<0.001, ****P<0.0001 by one-way ANOVA with Tukey correction for multiple comparisons. **(A-H)** Data representative of ≥3 independent experiments.

As mutant CD19^+^B220^−^ cells had similar IgM and CD5 profiles to what would be expected of wildtype B1 cells, we wanted to test whether these cells were indeed and expanded population of B1 cells. A tSNE analysis of total B cells in the spleen however, showed that mutant B cells clustered independently from other B cell populations (Fig. 3G). As expected, this analysis also showed an absence of the majority of normal B cells in *Pax5*^*Y351*/Y351\**^ mice (Fig. 3G). Strikingly, when wildtype and mutant cells were overlayed, *Pax5*^*Y351*/Y351\**^ CD19^+^B220^−^ cells (in red) grouped most closely to wildtype B1 (dark blue) and MZB cells (light blue) but did not overlay with either of these populations (Fig. 3H). This suggests that CD19^+^B220^−^ cells are similar to, but also distinct from B1 and MZB cells. To validate this, splenocytes were also stained for CD43 which is highly expressed on B1 cells, and for CD9 and CD1d, both of which are highly expressed on MZBs. Whilst CD43 was very lowly expressed on mutant CD19^+^B220^−^ cells compared to wildtype B1 cells (Fig. S4B), CD1d and CD9 were upregulated compared to wildtype MZB cells (Fig. S4C). This provides further evidence suggesting that mutant CD19^+^B220^−^ cells are neither B1 nor MZB cells but are a population that are unique to *Pax5*^*Y351*/Y351\**^ mice.

To further investigate the function of the truncated the PAX5 protein and the origin of the CD19^+^B220^−^ cells, we analyzed hematopoietic cell development in the bone marrow as well as expression of key PAX5 targets on early-stage B cells by flow cytometry. *Pax5*^*Y351*/Y351\**^ mice had normal numbers of lineage-negative hematopoietic stem cells (HSCs), multipotent progenitors (MPPs), common lymphoid progenitors (CLPs), and B cell primed lymphoid progenitors (BLPs) (Fig. S5, A and B). This suggests that the *Pax5*^*Y351\**^ mutation does not affect the development of early progenitors and that the abnormal CD19^+^B220^−^ cells originate from cells within the B cell lineage. Furthermore, a fraction of CD19^+^B220^−^ cells expressed μIgH, slightly increased to what is observed in wild-type pro-B cells, while the pro-B cells present in *Pax5*^*Y351*/Y351\**^ mice showed reduced expression of intracellular IgM heavy chain (Fig. 5A). To compare their likeness to pro-B cells, we used flow cytometry to analyze the expression of various direct targets of PAX5 to observe any changes that may result from the expression of truncated PAX5. This showed normal expression of most genes analyzed, with the exception of *Ly6e* (Sca-1) (Fig. 4B), where expression was significantly increased compared to wildtype suggesting dysregulation of specific target genes that are normally down-regulated by PAX5. Furthermore, the CD19^+^B220^−^ cells maintained Flt3 expression similar to CLPs in wildtype animals (Fig 4C). Together, these results suggest a failure to completely switch off B cell-inappropriate genes despite normal activation of B cell-appropriate genes both in pro-B cells and in the abnormal CD19^+^B220^−^ cells. As PAX5 expression is induced in pro-B cells (*15, 30, 31*), this data argues that CD19^+^B220^−^ cells are derived from pro-B cells that have failed to upregulate transcriptional programs necessary to completely commit to the B cell lineage.

**Fig. 4.**
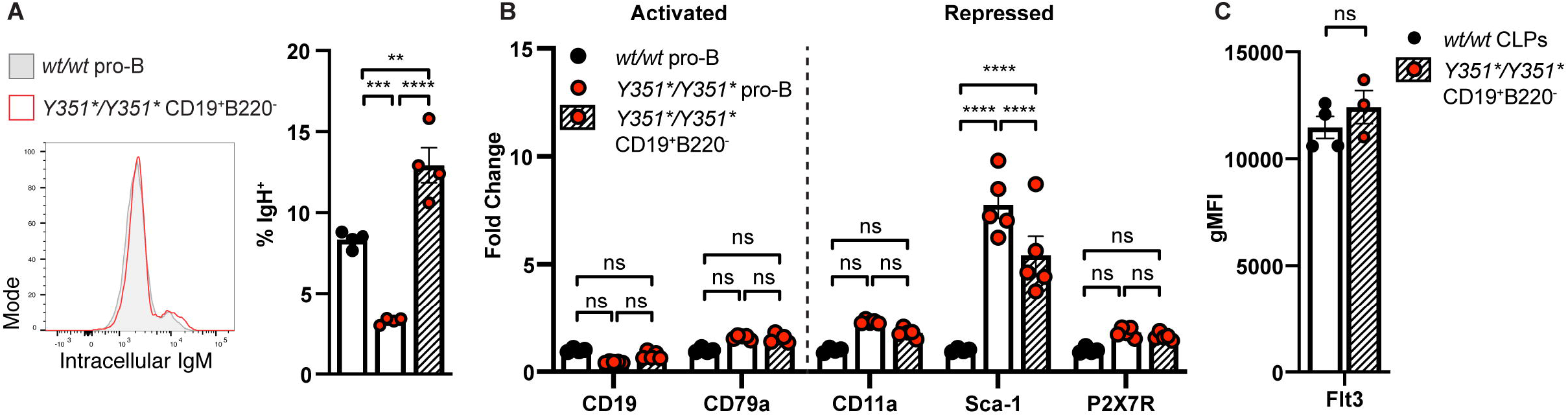
*Pax5*^*Y351*/Y351\**^ CD19^+^B220^−^ B cells and pro-B cells fail to downregulate Flt3 and Sca-1 despite residual PAX5 activity. **(A)** Representative histogram of intracellular IgM expression (μIgH) in *Pax5*^*wt/wt*^ pro-B cells (grey shaded) and *Pax5*^*Y351*/Y351\**^ CD19^+^B220^−^ cells (red line), determined by flow cytometry. Frequency of cells expressing intracellular IgM in *Pax5*^*wt/wt*^ pro-B cells, *Pax5*^*Y351*/Y351\**^ pro-B cells, and *Pax5*^*Y351*/Y351\**^ CD19^+^B220^−^ cells in the bone marrow determined by flow cytometry. **(B)** Fold change of expression (gMFI) of PAX5 activated and PAX5 repressed genes in *Pax5*^*wt/wt*^ pro-B cells, *Pax5*^*Y351*/Y351\**^ pro-B cells, and *Pax5*^*Y351*/Y351\**^ CD19^+^B220^−^ cells in the bone marrow, determined by flow cytometry. (C) Expression (gMFI) of Flt3 on *Pax5*^*wt/wt*^ common lymphoid progenitors (CLPs) and on *Pax5*^*Y351*/Y351\**^ CD19^+^B220^−^ cells in the bone marrow. **(A-C)** Bars show average per group ± SEM with individual mice indicated by circles. Asterisks indicate significance: ns P≥0.05, *P<0.05, **P<0.01, ***P<0.001, ****P<0.0001 by two-way ANOVA with Tukey correction for multiple comparisons. **(A)** Data representative of ≥3 independent experiments. **(B, C)** Data obtained from one experiment.

**Fig. 5.**
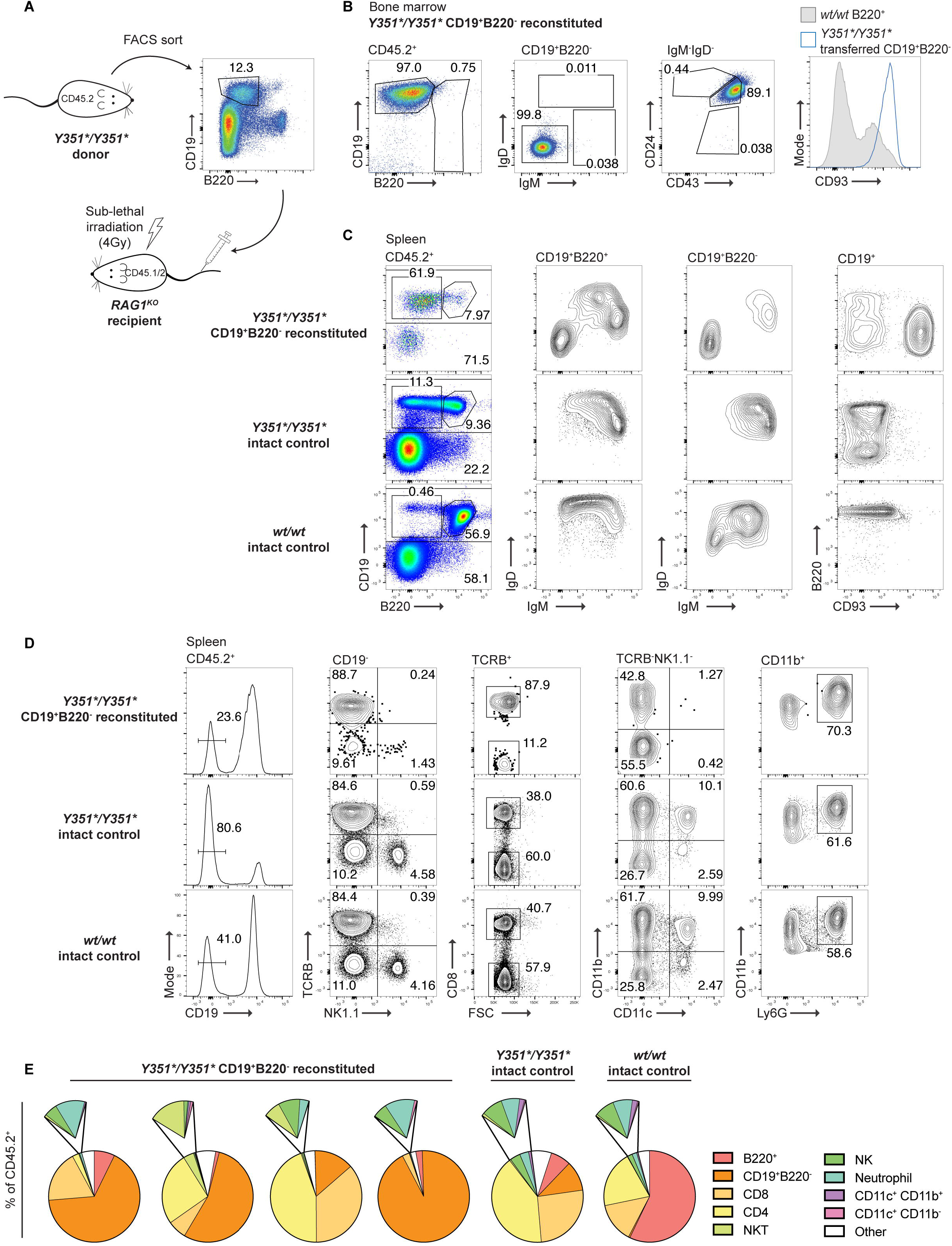
Abnormal CD19^+^B220^−^ B cells have self-renewing and multipotent capacities. **(A)** Schematic of *Rag1*^−/−^ reconstitution with *Pax5*^*Y351*/Y351\**^ cells. CD19^+^B220^−^ cells were isolated from the bone marrow of CD45.2 *Pax5*^*Y351*/Y351\**^ mice by FACS isolation. 5×10^6^ CD19^+^B220^−^ cells were transferred into sub-lethally irradiated (4Gy) CD45.1/2 *Rag1*^−/−^ mice by tail IV injection. The bone marrow and spleens of recipient mice were analyzed 8 weeks post transfer. **(B)** Representative FACS plots of donor-derived (CD45.2^+^) cells in the bone marrow of reconstituted *Rag1*^−/−^ mice showing maintenance of identity post transfer. **(C)** Representative FACS plots of donor-derived (CD45.2^+^) B cells in the spleen of reconstituted *Rag1*^−/−^ mice showing IgM and IgD expression on B220^+^ and B220^−^ B cells, pre-gated on CD19^+^, compared with intact *Pax5*^*Y351*/Y351\**^ and *Pax5*^*wt/wt*^ controls. **(D)** Representative FACS plots of donor-derived non-B cells in the spleen of reconstituted *Rag1*^−/−^ mice compared to intact *Pax5*^*Y351*/Y351\**^ and *Pax5*^*wt/wt*^ controls. **(E)** Frequency of lymphoid and myeloid cell subsets gated from total CD45.2^+^ donor-derived cells. Each pie chart represents one mouse with each cell type represented in different colors as indicated. **(B-D)** Reconstituted *Rag1*^−/−^ n=4, intact *Pax5*^*Y351*/Y351\**^ controls n=1, intact *Pax5*^*wt/wt*^ controls n=1. Data representative of one experiment.

To determine the developmental potential of CD19^+^B220^−^ B cells, we FACS sorted CD19^+^B220^−^IgM^−^CD93^+^CD45.2^+^ cells from *Pax5*^*Y351*/Y351\**^ bone marrow and adoptively transferred them into sub-lethally irradiated CD45.1/2 *Rag1*^−/−^ hosts (Fig. 5A). 8 weeks post-transfer, the bone marrow of recipient mice showed that almost all of the CD45.2^+^ cells had maintained their original CD19^+^B220^−^IgM^−^CD93^+^ identity (Fig. 5B), indicating that the mutant cells were able to engraft and self-renew. “Normal” CD45.2^+^B220^+^ B cells were unable to be identified in the recipients’ bone marrow (Fig. 5B). In contrast, CD45.2^+^B220^+^ cells could be found in the spleen of recipient mice and were either IgM^−^IgD^−^ or IgM^+^IgD^low^ (Fig. 5C). This is in comparison to intact *Pax5*^*Y351*/Y351\**^ mice where all B220^+^ B cells were IgM^+^IgD^low^ (Fig. 5C). Additionally, the abnormal population of CD19^+^B220^−^ cells present in the spleen of *Pax5*^*Y351*/Y351\**^ mice could also be found in the recipient spleens. But in contrast to the cells found in intact mice, the CD19^+^B220^−^ cells in recipient mice expressed high levels of the immature cell marker CD93 (Fig. 5C) suggesting that the transferred cells were not able to undergo the normal maturation in the spleen. In agreement, the absence of B220^+^ B cells in the bone marrow of reconstituted recipients also suggests that CD19^+^B220^−^ B cells are unable to progress through normal B220^+^ developmental stages within the bone marrow and spleen and that B220 expression is obtained at some point in the periphery.

### Subhead 4: Correct maintenance of PAX5 expression prevents plasticity in B cells

*Pax5*^−/−^ pro-B cells are not committed to the B cell lineage and are therefore able to de-differentiate into other hematopoietic cell lineages (*15*). Furthermore, deleting PAX5 from mature B cells reverts their commitment to the B cell lineage and allows them to de-differentiate into T and NK cells (*20*). To see if the PAX5^Y351*^ CD19^+^B220^−^ B cells also maintained a hematopoietic plasticity, the spleens of CD19^+^B220^−^ reconstituted *Rag1*^−/−^ mice were analyzed for the presence of different leukocyte populations. Surprisingly, not only could *Pax5*^*Y351*/Y351\**^ CD19^+^B220^−^ B cells differentiate into both CD4^+^ and CD8^+^ T cells (TCRβ^+^), as well as NK cells (NK1.1^+^), but they could also differentiate into myeloid cells, including dendritic cells (CD11b^+^CD11c^+^) and neutrophils (CD11b^+^Ly6G^+^) (Fig. 5D). Results between recipients were variable, but nevertheless CD45.2^+^ cells of both lymphoid and myeloid origin were able to be identified in all recipients (Fig. 5E). This suggests that the CD19^+^B220^−^ mutant bone marrow cells have a broad multi-lineage potential and are therefore unlike *Pax5*^−/−^ pro-B cells as they maintain an ability to develop into mature B cells. Importantly, this is the first time that it has been shown that a B cell still expressing PAX5 is able to be converted to a cell of a different lineage through *in-vivo* mechanisms.

### Subhead 5: Y351* mutation causes an overall reduction in PAX5 function

Evidence presented thus far demonstrates that PAX5^Y351*^ does not behave as a null allele. The presence of B cells in *Pax5*^*Y351*/Y351\**^ mice suggests that some transcriptional activation and repression by PAX5 is retained and allows for the establishment of B cell identity in select clones. To understand how the *Pax5*^*Y351\**^ mutation was affecting protein function, we performed RNAseq of CD19^+^B220^−^ cells from the bone marrow of *Pax5*^*Y351*/Y351\**^ mice and compared this to B220^+^ pro-B cells from *Pax5*^*Y351*/Y351\**^ and *Pax5*^*wt/wt*^ mice.

By filtering down to genes that are either directly activated or repressed by PAX5 (*7*) the exact effect of the Y351* mutation on PAX5 function was determined. Genes that are activated by PAX5 were generally decreased in both *Pax5*^*Y351*/Y351\**^ CD19^+^B220^−^ cells and pro-B cells compared to wildtype pro-B cells (Fig. 6A and Table S1). Conversely, genes that are repressed by PAX5 were more generally increased in mutant cells compared to wildtype pro-B cells (Fig. 6A and Table S1). This suggests that the Y351* mutation results in an overall reduction in PAX5 activity. A gene set enrichment analysis (GSEA) of genes positively regulated by PAX5, but not necessarily direct DNA binding targets of PAX5 (*7*) showed that wildtype pro-B cells are significantly enriched for PAX5 activated genes compared mutant pro-B cells (Fig. 6B). GSEA of genes negatively regulated by PAX5 failed to show any statistically significance difference in expression between wildtype and mutant pro-B cells (Fig. 6B), possibly because the gene set is relatively small to be included in a traditional GSEA. To confirm that this was an effect specific to PAX5-regulated genes rather than random variation within the dataset, we compared the expression of three independent housekeeping gene sets (Table S2) across the three cell types. The null hypothesis was that genes unrelated to PAX5 would have a comparable number of reads in both mutant and wildtype B cells. Gene expression was normalized to the maximum number of reads for each specific gene across all cell types, and the average gene expression per cell type was determined. In agreement with the null hypothesis, we found that *Pax5*^*Y351*/Y351\**^ pro-B and CD19^+^B220^−^ cells had no statistically significant difference in the average expression level of the three housekeeping gene sets compared to wildtype pro-B cells (Fig. S6). Furthermore, when the same analysis was applied to PAX5-regulated gene sets, significant differences were observed in the overall expression of both PAX5-activated and PAX5-repressed in *Pax5*^*Y351*/Y351\**^ pro-B and CD19^+^B220^−^ compared to wildtype pro-B cells (Fig. S6). This demonstrates that the Y351* mutation is affecting the efficacy of gene regulation performed by PAX5 on a population level. Interestingly the expression of PAX5-regulated genes was comparable in mutant pro-B and CD19^+^B220^−^ cells (Fig. S6), indicating a similar effect of the Y351* mutation on the PAX5-transcriptome in these two cell types.

**Fig. 6.**
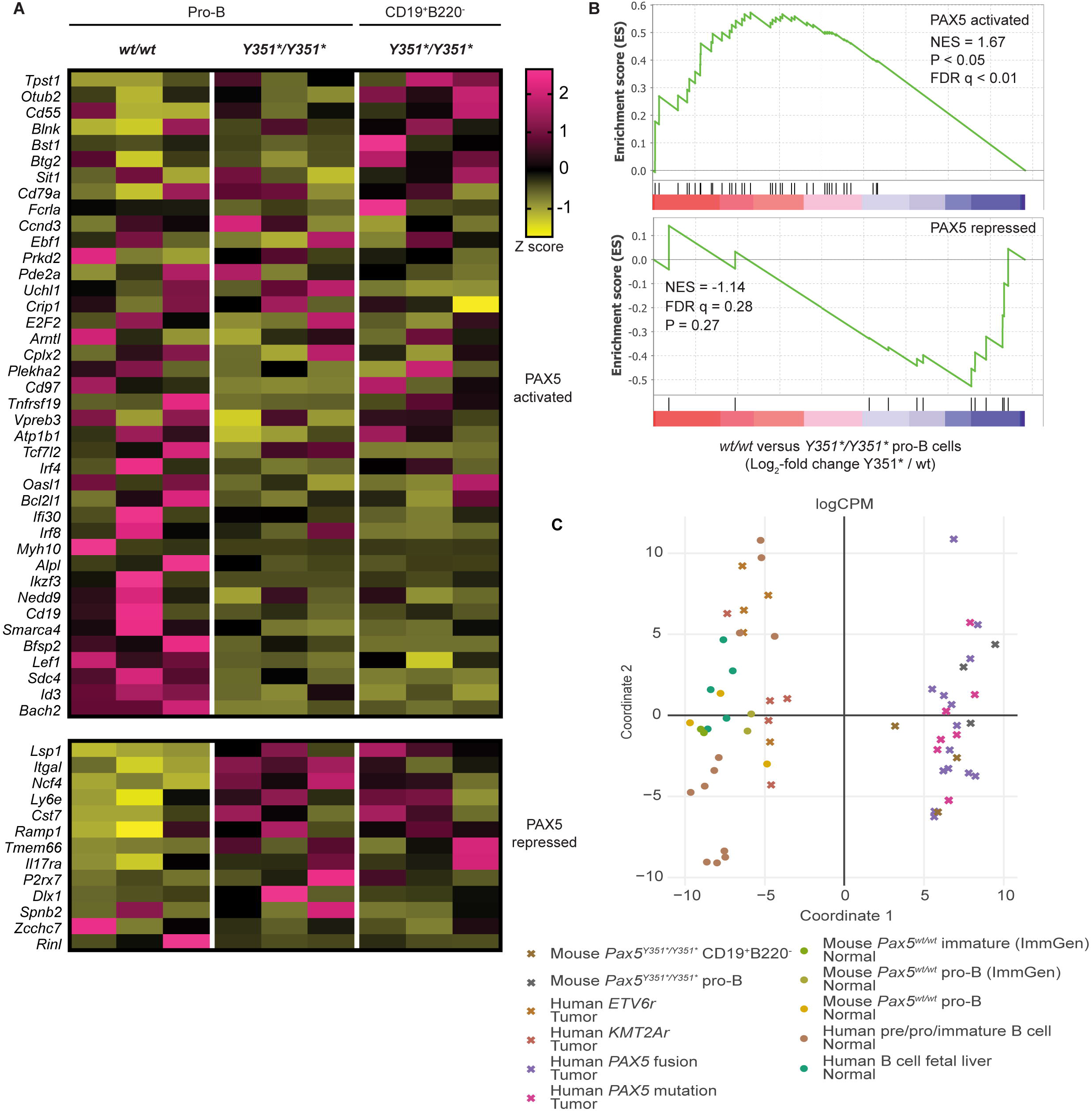
PAX5^Y351*^ causes an overall loss of function with reduced activation and repression of PAX5 target genes. **(A)** RNAseq heatmap showing the Z score of genes directly bound by PAX5 in wildtype pro-B cells (*7*) compared to *Pax5*^*Y351*/Y351\**^ pro-B and CD19^+^B220^−^ cells. **(B)** Gene set enrichment analysis of PAX5 activated (top) and PAX5 repressed (bottom) genes in *Pax5*^*wt/wt*^ pro-B cells compared to *Pax5*^*Y351*/Y351\**^ pro-B cells using the Log2 fold change of mutant compared to wildtype. NES: normalized enrichment score, FDR: false discovery rate. **(A-B)** 50 cells per population FACS isolated with 4 technical repeats pooled together post sequencing, *wt/wt* n=3, *Y351*/Y351\** n=3. **(C)** Multidimensional scaling (MDS) plot based on the 48 differentially expressed genes shown in Figure 6 showing the clustering of pro-B cells and CD19^+^B220^−^ cells from *Pax5*^*Y351*/Y351\**^ mice with B-ALL cells from humans with mutations in PAX5, whilst B-ALL cells driven by other genetic changes cluster with normal human and mouse pro-B cells. **(A-C)** Data obtained from one experiment.

We then compared how the transcriptome of PAX5-regulated genes in *Pax5*^*Y351*/Y351\**^ mice compared with tumors from patients with *PAX5* fusion or *PAX5* point mutations (Fig. 6C and Table S3). As controls, we compared tumors from patients with unrelated mutations in *ETV6r* or *KMT2Ar* and well as data sets obtained for human pre, pro, and immature B cells and human fetal liver B cells. To control for differences in mouse-to-human regulation, we included our own *Pax5*^*wt/wt*^ pro-B cells as well as additional data from wildtype pro-B and immature B cells from the ImmGen database. Multidimensional scaling (MDS) of PAX5-regulated genes showed clustering of B cells with PAX5 mutations from both mouse and human datasets (Fig. 6C). Wildtype B cells and tumors with *ETV6r* or *KMT2Ar* mutations clustered together demonstrating validity of the dataset and likeness of human and mouse regulation of PAX5-target genes. Together this shows how murine PAX5 acts similarly in health and disease to human PAX5 and provides evidence of the similarities between dysregulation in *Pax5* mutant B cells in mice, and *PAX5-*driven B-ALL in humans.

### Subhead 6: Dysregulation of PAX5 induces oncogenic changes and the development of precursor B cell leukemia

Strikingly, when *Pax5*^*Y351*/Y351\**^ mice were aged, they developed gross splenomegaly and lymphadenopathy (Fig. 7A) causing acute illness with a 100% penetrance and a median survival of 239 days (Fig. 7B). Female mice had an earlier median onset of disease compared with male littermates at 197 compared to 247 days respectively (Fig. S7). Affected mice could be readily identified by an expansion of large, CD19^+^B220^−^CD93^+^IgM^−^ cells in the blood (Fig. 7C). Onset of disease was very quick, with a small increase in the frequency of tumor cells first detectable in the blood 2-4 weeks before overt disease progression resulted in euthanasia of the affected mouse. Lymph node enlargement however, was only obvious a couple of days prior to euthanasia. *Pax5*^*wt/Y351\**^ mice also developed similar clinical symptoms, except with a reduced frequency and a greater median age of onset (744 days) compared to *Pax5*^*Y351*/Y351\**^ mice (Fig. 7B). Except for some variation in B220 expression between mice, cells isolated from affected *Pax5*^*Y351*/Y351\**^ mice showed a consistent phenotype of CD19^+^CD93^+^IgM^−^ (Fig. 7C). At the time of takedown, the lymph nodes, spleen, and bone marrow from affected mice showed almost a complete dominance of these cells. The expanded cells were also negative for intracellular IgM staining (Fig. S8), indicating the μIgH had not yet been expressed therefore suggesting a precursor B cell origin.

**Fig. 7.**
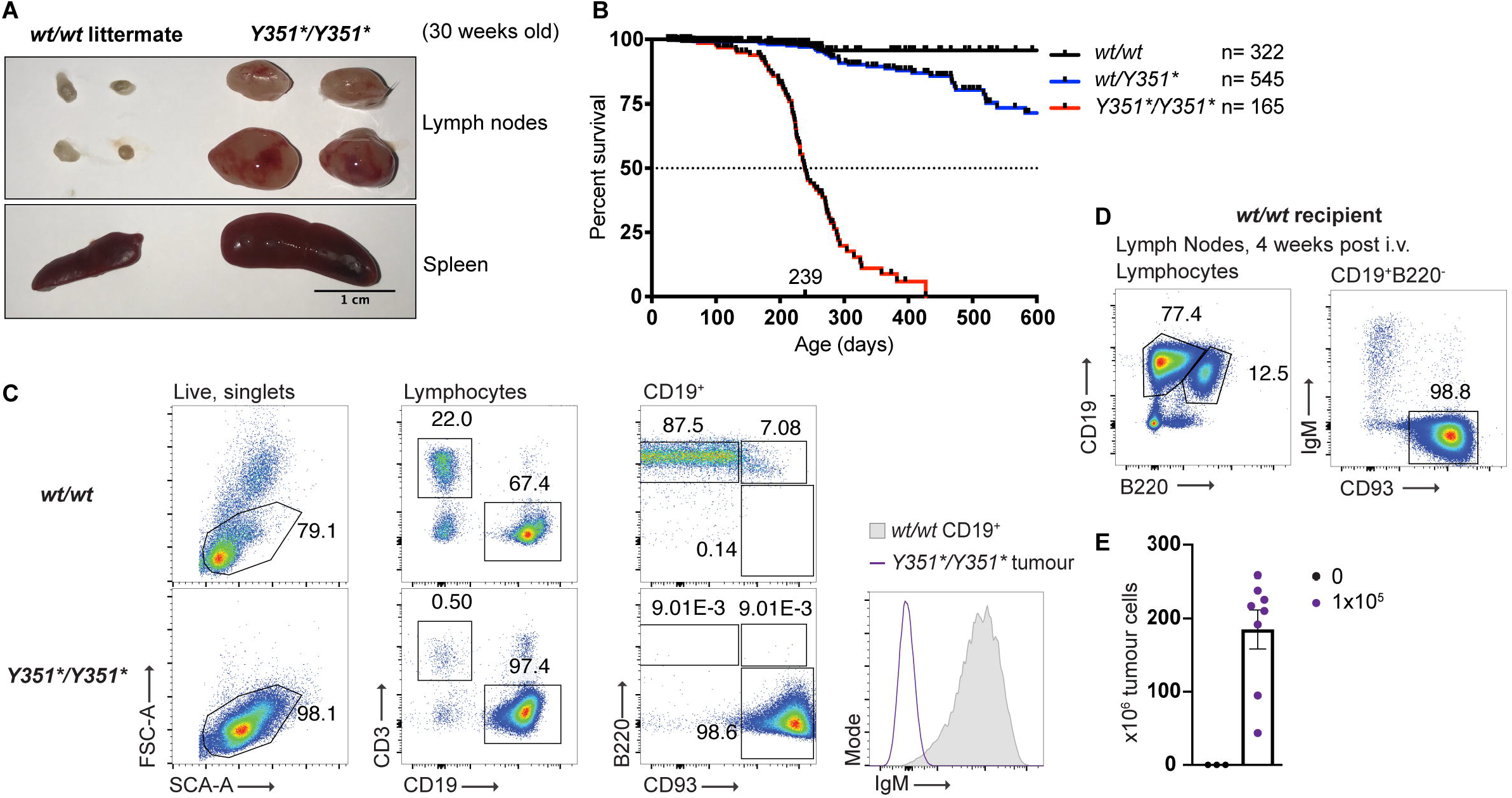
*Pax5*^*Y351*/Y351\**^ mice spontaneously develop B cell acute lymphoblastic leukemia with 100% penetrance. **(A)** Lymph nodes (x2 brachial & x2 inguinal) and spleen from aged *Pax5*^*Y351*/Y351\**^ and Pax5^*wt/wt*^ littermates. Scale bar=1cm. **(B)** Kaplan-Meier survival curve showing death by confirmed tumor development or death due to unknown or undocumented reasons. Death by unrelated reasons (e.g. malocclusion or fight wounds) were censored. Median survival for homozygous mice is 239 days. **(C)** Representative FACS plots of peripheral blood from an aged *Pax5*^*Y351*/Y351\**^ mouse with lymphadenopathy, demonstrating a homogenous expansion of an immature B cell population in comparison to a wildtype littermate. Data representative of ≥50 homozygous mice. **(D)** 1×10^5^ tumor cells from the lymph nodes of affected mice were adoptively transferred into wildtype mice. FACS plots show lymphocytes isolated from the lymph nodes of a recipient mouse, 3 weeks post transfer. **(E)** Total number (x10^6^) of tumor cells isolated from the lymph nodes (x2 brachial and x2 inguinal) of recipient wildtype mice three weeks post transfer. **(D & E)** Data representative of 2 independent experiments.

To confirm the malignant potential of these expanded cells, 1×10^5^ cells from enlarged lymph nodes of *Pax5*^*Y351*/Y351\**^ mice were adoptively transferred either as a total cell suspension or as an isolated FACS sorted population, into wildtype C57BL/6 mice. Within 3-4 weeks, recipient mice had to be euthanized due to loss of condition and severe lymphadenopathy, similar to what was observed in aged intact *Pax5*^*Y351/Y351\**^ mice. Transferred cells maintained their CD19^+^B220^−^IgM^−^CD93^+^ phenotype (Fig. 7D) and could be identified in large numbers in the lymph nodes of recipient mice (Fig. 7E). The precursor origin of these malignant B cells and the acute onset of disease along with the known tumor-suppressor role of PAX5 (*23*), suggests that the PAX5^Y351*^ mutation drives the spontaneous development of B-ALL in mice. As the tumor cells isolated from *Pax5*^*Y351*/Y351\**^ mice had a very similar phenotype to the multipotent CD19^+^B220^−^ cells present in the bone marrow of young homozygous mice, we hypothesized that the abnormal CD19^+^B220^−^ bone marrow cells were a pre-malignant precursor to the tumor cells identified in aged homozygous mice. In support of this, one otherwise unmanipulated mouse received sorted, abnormal CD19^+^B220^−^ bone marrow cells from a young *Pax5*^*Y351*/Y351\**^ mouse and one year after transfer developed gross lymphadenopathy and severe loss of condition due to an accumulation of donor-derived CD19^+^B220^−^ cells. This corresponds with data demonstrating similarities between *Pax5*^*Y351*/Y351\**^ CD19^+^B220^−^ cells and human *PAX5*-driven B-ALL (Fig. 6C), suggesting a potential origin of B-ALL in *Pax5*^*Y351*/Y351\**^ mice.

### Subhead 7: JAK3 and PTPN11 cooperate with PAX5 to induce malignant transformation

Given the age in which *Pax5*^*Y351*/Y351\**^ mice develop tumors, it is likely that with *Pax5*^*Y351\**^ as the driver mutation, precursor B cells develop secondary mutations that induce malignant transformation and uncontrolled proliferation. As such, we performed whole exome sequencing on sorted tumor cells isolated from 14 different *Pax5*^*Y351*/Y351\**^ mice. This identified 113 different mutations across the 14 tumors with a range of 1-20 and an average of 10.5 mutations per tumor (Fig. 8A and Table S4). Quite strikingly, 6 out of 14 tumors had the exact same SNP in *Jak3* (R653H) with a further 2 tumors identified to also carry different non-synonymous *Jak3* mutations (T844M and A568V) (Fig 8B and Table S4). Additionally, 4 different SNPs in *Ptpn11* (S502P, S502L, N58Y, and E69K) were identified in 5/14 tumors. Concurrent mutations in both *Jak3* and *Ptpn11* were present in 4/14 tumors (Fig. 8A). A non-synonymous SNP in *Zmiz2* (G575A) was identified in 5/14 tumors but was later identified to have arisen from a spontaneous germline mutation within the colony so was removed from the analysis (Table S4).

**Fig. 8.**
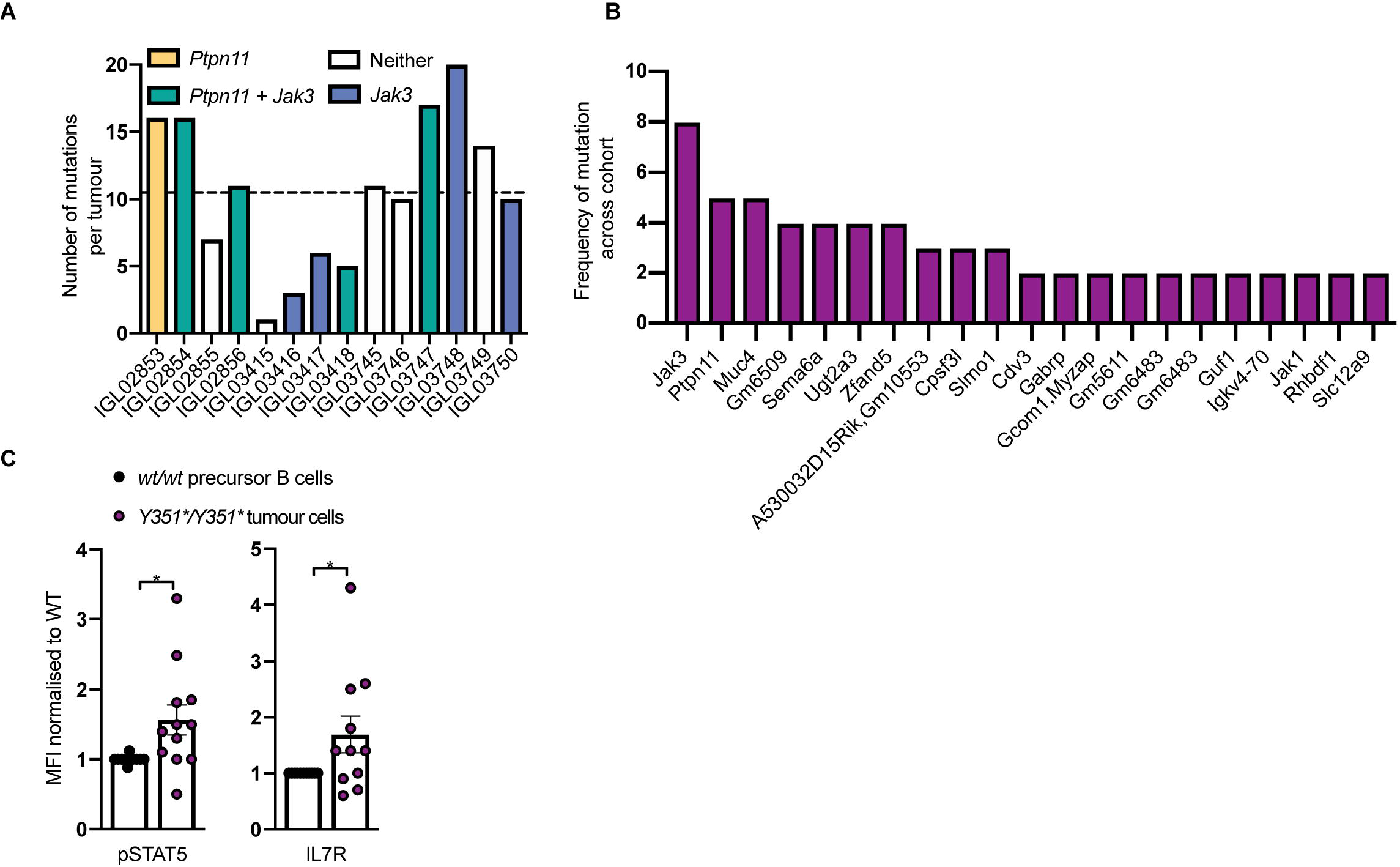
PAX5^Y351*^ acts synergistically with secondary *Jak3*, and *Ptpn11* mutations leading to constitutively activated IL-7 and STAT5 signaling. **(A)** Total number of mutations per tumor from *Pax5*^*Y351*/Y351\**^ lymph nodes by whole exome sequencing. Each bar represents bulk tumor cells isolated from individual mice. Colors of bars indicate combinations of mutation in *Jak3* and/or *Ptpn11* as indicated. White bars represent tumors where no mutations in either *Jak3* or *Ptpn11* were identified. **(B)** Occurrence of mutations in indicated genes across the cohort of sequenced tumors. Results filtered to include genes where mutations were identified in at least two tumors; n=14 tumors. **(C)** Expression of pSTAT5 and IL-7R by flowcytometry in *Pax5*^*Y351*/Y351\**^ tumor cells, normalized to wildtype precursor B cells (B220^+^IgM^−^IgD^−^). gMFI from each tumor was normalized to either a wildtype littermate or an age-matched control within the same experiment and data pooled together post analysis. Bars show average per group ± SEM with individual mice indicated by circles. Asterisks indicate significance: *P<0.05 by students two-tailed T test.

The R653H *Jak3* mutation is a known oncogenic mutation, causing constitutive activation of STAT5 through IL-7 signaling (*32*). Accordingly, resting *Pax5*^*Y351*/Y351\**^ tumor cells had on average increased expression of both pSTAT5 (Y694) and IL-7R compared to precursor B cells from wildtype bone marrow (Fig. 8C), thus confirming increased steady-state activation of this pathway.

## Discussion

Thus far, *in vitro* studies have identified four different functional domains of PAX5, but there have been no studies that have translated these findings *in vivo.* The Y351* mutation is unique in that it provides mechanistic insight not only into the *in vivo* function of the PAX5 inhibition domain, but also into the final 8 amino acids of the transactivation domain. Importantly, it is the first demonstration of an *in vivo* model that stably expresses a truncated version of PAX5 (Fig. 1D). We show that the truncated protein has dramatic effects on B cell lineage commitment and causes the spontaneous development of B-ALL. A key question arising from our study is how B cells can develop in the absence of full length PAX5, as previous studies have shown that PAX5 is vital for ensuring maintenance of B cell identity and without which, both pro-B (*15*) and mature B cells (*20*) become uncommitted to the B lineage, completely preventing the ongoing development of circulating B cells. In contrast, *Pax5*^*Y351*/Y351\**^ mice have low frequencies of circulating mature B cells, demonstrating that progression through normal B cell development is not a strict binary “on or off” decision that is determined by PAX5 and that reduced PAX5 functionality in a small proportion of pro-B cells can still result in the development of mature B cells. The absence of wildtype PAX5 however, leads to the expansion of a CD19^+^B220^−^ population in the bone marrow. Two possible avenues exist for the development of these abnormal B220^−^ B cells; either they are derived from HSCs and develop concurrently with normal lymphopoiesis in the bone marrow, or they are derived from pro-B cells that were unable to progress through to B220^+^ pre-B cells due to attenuated PAX5 signaling. As PAX5 expression is first induced in pro-B cells (*11, 14, 33*), it is unlikely that a mutation in *Pax5* would influence events prior to its upregulation and it is therefore more probable that CD19^+^B220^−^ cells are a product of failed pro-B cell development. It will be important for further studies to determine whether the immature IgM^+^ B cells that leave the bone marrow derive from the few pro-B cells found in *Pax5*^*Y351*/Y351\**^ mice, or if they develop from the unusual CD19^+^B220^−^ cells found in these animals. Transfer of these CD19^+^B220^−^ cells into sub-lethally irradiated *Rag1*^−/−^ hosts however, results in only very few B220^+^IgM^+^ cells (Fig. 5). This suggests that B cell differentiation in intact mice will most likely occur from the B220^+^ pro-B cells and that the inhibition domain, and possibly the transactivation domain, are crucial for the maintenance of the normal B cell program.

Our RNA-sequencing results and flow-cytometric determination of protein expression of key PAX5 target genes have revealed a surprisingly variable role of the inhibition domain and parts of the transactivation domain. Some well-validated PAX5 target genes such as *Cd19,* showed an almost normal upregulation *in vivo* (Fig. 1B and Fig. 4B). This contrasts with previous *in vitro* studies that found that the whole transactivation domain was crucial for the normal expression of CD19 (*5, 34, 35*). Other genes showed a normal down-regulation even in the absence of full-length PAX5, while *Ly6e* (Sca-1) showed a complete failure to downregulate in pro-B cells (Fig. 4B) and the CD19^+^B220^−^ cells maintained high expression of Flt3 similar to levels observed in CLPs (Fig. 4C). This variation in the response of well-validated PAX5 target genes indicates differential requirements of PAX5 functional domains in either the activation or repression of individual genes, rather than a global loss-of-function. This further emphasizes the difference between *Pax5*^*Y351*/Y351\**^ and *Pax5*^−/−^ mice and highlights a previously unknown complexity in PAX5 function.

The observation that the CD19^+^B220^−^ cells found in the bone marrow of *Pax5*^*Y351*/Y351*^ mice can transdifferentiate into different hematopoietic lineages (Fig. 5D), despite expressing comparable PAX5 levels to wild-type pro-B cells (Fig. S4A), suggests a key role for the inhibition domain in shutting down multi-lineage potential. Similarly to *Pax5*^*−/−*^ pro-B cells (*15*) and mature B cells (*20*) that have lost PAX5 expression, *Pax5*^*Y351*/Y351\**^ CD19^+^B220^−^ cells maintain a multipotent, self-renewing phenotype, with a broad lympho-myeloid potential (Fig. 5). Despite a clear loss of the PAX5-driven B cell program in some cells, unlike in *Pax5*^−/−^ mice not all *Pax5*^*Y351*/Y351\**^ progenitor B cells have completely lost their ability to differentiate into mature B cells. Future studies will be required to determine if the multilineage potential and maintenance of B cell potential reside within the same cell or reflect a further loss of PAX5 function in a subset of the CD19^+^B220^−^ cells found in *Pax5*^*Y351*/Y351\**^ mice. In addition to their multilineage potential, a sizable fraction of cells found in the periphery of mice 8 weeks after transfer of B220^−^CD19^+^ cells show a phenotype very similar to the B-ALL developing in intact *Pax5*^*Y351*/Y351\**^ mice, strongly suggesting that the B220^−^CD19^+^ B cells found in these mice are the origin of the B-ALL.

Hypomorphic *PAX5* mutations are the most common driver of pediatric B-ALL (*21, 23*), yet the effects of reduced, as opposed to absent, PAX5 function on B cells have remained elusive. *Pax5*^−/−^ mice die prematurely (*14*) and hence studying the effects of PAX5-deficiency has been difficult. It has also not been possible to study the effects of PAX5-haploinsufficiency on tumor development, as *Pax5*^*+/−*^ mice develop no tumors unless crossed with mice that have constitutively active expression of known oncogenic proteins (*27*). Here, we present the first in-depth analysis of spontaneous PAX5-driven B-ALL in a germline mouse-model. Additionally, we present the first direct comparison of murine *Pax5* mutant B cells with *PAX5-* driven B-ALL in humans, validating the use of *Pax5*^*Y351*/Y351\**^ mice as a model for human B-ALL. Consistent with previous work in both murine and human B-ALL, PAX5^Y351*^-driven tumors have common secondary mutations in *Jak3* and *Ptpn11,* (Fig. 8, A and B) (*23, 26, 32*) that facilitate transcription of survival and growth factors. The *Pax5*^*Y351*/Y351\**^ tumors harbor the same secondary mutations discovered in human B-ALL and hence provide an accurate tool for studying PAX5-driven B-ALL *in vivo.* Additionally, the CD19^+^B220^−^ cells found in pre-malignant *Pax5*^*Y351*/Y351\**^ mice have a very similar phenotype to *Pax5*^*Y351*/Y351\**^ tumor cells, which suggests that they are a precursor to tumor cells.

The *Pax5*^*Y351*/Y351\**^ mice provide novel insight into the regulatory mechanisms of PAX5. They demonstrate that full-length PAX5 is absolutely required for the development, maintenance, and commitment of B cells. Removing the inhibitory domain of PAX5 is sufficient to attenuate B cell development and reroute precursor B cells through an alternate fate, producing non-committed, tumorigenic B cells. This leads to an accumulation of lineage-promiscuous B cells that eventually develop secondary oncogenic lesions, ultimately causing malignant expansion and escape from the bone marrow. In some precursor cells, B cell development is able to occur normally and resulting mature B cells appear phenotypically normal with the exception of downregulated IgD, CD23, and CD21/35. The *Pax5*^*Y351*\*^ mutation confirms previously known roles of the inhibition domain, but challenges data regarding the transcription of *Cd19* by truncated PAX5 protein (*5, 34, 35*). Importantly, the *Pax5*^*Y351*/Y351\**^ mice provide an invaluable asset to study the effects of PAX5 mutations on the development and treatment of B-ALL in mammals. Analyzing the outcomes of novel therapies on PAX5^Y351*^ tumors *in vivo* will be essential for the future development of effective treatment regimens for PAX5-driven B-ALL.

## Materials and Methods

### Study Design

The *Pax5*^*Y351\**^ mice were originally identified through an *N*-ethyl-*N*-nitrosourea (ENU)-screening project aimed to identify novel regulators of B cell development. Subsequently, this project was designed to characterize the role of the PAX5 inhibition domain on B cell development. This is the first *in vivo Pax5* mutant that stably expresses PAX5 and survives well into adulthood. We took advantage of this model by designing experiments that characterize all stages of B cell development throughout primary and secondary lymphoid organs. At least 3 mice per group within each independent experiment were used to determine statistical significance. The number of independent repeats per experiment are included in the figure legends but data from only one representative experiment is shown. All repeats demonstrated the same trend as those shown in the figures. 3 data points were excluded in Fig. 1A as mice had obviously developed tumors that were causing a substantial increase in CD19^+^ B cell frequency (48.1%, 62.9%, and 97.1% respectively) and therefore skewing the group average. No other data points were excluded.

### Mice

The *Pax5*^*Y351\**^ mutant strain was identified by flow cytometric screening of blood lymphocytes in third generation offspring of C57BL/6J mice treated with 3 × 90-100 mg/kg ENU as described previously (*36*). Variant calling from whole exome sequencing data was performed as previously described (*37*). All mice were generated on a C57BL/6J background and subsequently backcrossed to a C57BL/6NCrl for a minimum of 10 generations. For all mouse experiments, littermate or age-matched controls were used. Mice were housed in specific pathogen–free conditions at the Australian National University Bioscience Research Services facility, and all animal procedures were approved by the Australian National University Animal Ethics and Experimentation Committee on protocols A2011/46, A2014/62 and A2017/54.

### Flow Cytometry

Analysis of peripheral blood was performed on 6-15 week old mice as previously described (*38*). For analysis of lymphocyte development, pre-malignant 8-15-week-old naïve mice were sacrificed through cervical dislocation or CO_2_ inhalation and organs collected in FACS buffer (1X PBS, 2.5% HI-BS, 0.1% NaN_3_). Bone marrow was flushed out from the femur and tibia of one hind leg per mouse using a 26G needle and filtered through a 70μm cell strainer. Single-cell suspensions for all other organs were mechanically disrupted and filtered through a 70μm cell strainer. Spleens were treated with red blood cell lysis buffer (8.99% w/v NH4Cl, 1% w/v KHCO_3_, 0.037% w/v EDTA, pH=7.3) and then washed with FACS buffer. Cells of the peritoneal cavity were obtained by injecting 3mL of FACS buffer into the peritoneum with a 23G needle and re-extracting ~2.5mL of fluid after palpation of the peritoneum. Samples were acquired using an LSR II, LSRFortessa™, or LSRFortessa™ X-20 (BD Bioscience) and analyzed using FlowJo software version 10 (FlowJo LLC). For intracellular staining, cells were treated with Foxp3 Transcription Factor Staining Buffer Set (eBioscience) according to the manufacturer’s instructions. For phosphoflow, cells were fixed with 1.5% paraformaldehyde for 10 minutes, washed thoroughly with PBS, and then permed with ice-cold methanol and stored at −20°C overnight before washing thoroughly with PBS and staining with fluorescently labelled antibodies.

### Western Blot

Cells were obtained from the bone marrow of pre-malignant 10-week-old mice as described above. B cells were isolated by labelling with αCD19-PE and then incubating with αPE microbeads (Miltenyi) and passed through a MACS LS column (Miltenyi) according to the manufacturer’s instructions. Lysate from 1×10^6^ positively sorted B cells were prepared in lysis buffer (50mM Tris [pH 7.4], 150mM NaCl, 2mM EDTA, 0.5% Triton X-100, and Halt protease and phosphatase inhibitor cocktail (Thermo Fisher Scientific)) by incubating for 30 minutes on ice. Lysate was then prepared with NuPAGE™ LDS Sample Buffer (4X) and Sample Reducing Agent (10X) (Invitrogen™) and boiled at 90°C for 5 minutes and frozen at −20°C for up to one month. Equal volumes of lysate were loaded onto NuPAGE 4-12% Bis-Tris 10 well gels (Invitrogen™) alongside Precision Plus Protein™ Kaleidoscope™ Prestained Protein Standard (Bio-Rad). Gels were run with NuPAGE™ MES SDS Running Buffer (20X) (Invitrogen™) for 35 minutes at 165V and transferred onto nitrocellulose membranes using the iBlot^®^ 7-Minute Blotting System (Thermo Fisher Scientific) according to the manufacturer’s instructions. Membranes were cut in half at approximately 75kD and blocked with 5% (w/v) skim milk in 0.1% Triton X-100 in PBS for 1 hour at room temperature. Membranes were then incubated with rat αmouse PAX5 (Biolegend) or rabbit αmouse VINCULIN (Cell Signaling Technology) diluted in 5% (w/v) skim milk in 0.1% TBST overnight at 4°C on a shaker. Membranes were then washed with 0.1% TBST and incubated with either αrat-HRP (Cell Signaling Technology) or αrabbit-HRP (Cell Signaling Technology) diluted 1:5000 in 5% w/v skim milk for 1 hour at room temperature. Membranes were washed again with 0.1% TBST and then developed with SuperSignal™ West Femto Maximum Sensitivity Substrate (Thermo Fisher Scientific) and viewed on a ChemiDoc MP Imaging System (Bio-Rad).

### FACS Cell Sorting

For adoptive transfer into sub-lethally irradiated mice, bone marrow cells from a 7-week-old *Pax5*^*Y351*/Y351\**^ mouse were collected in complete RPMI (2mM L-glutamine, 100U/mL penicillin, 100μg/mL streptomycin, 10% (v/v) FCS) and prepared as above. Cells were then incubated in 500μl of Fc Block in PBS for 30 minutes at 4°C, washed, and then stained in 500μl of antibody cocktail mix in PBS for 30 minutes at 4°C. 7AAD^−^CD19^+^B220^−^IgM^−^CD93^+^ lymphocytes were sorted on a FACS ARIA-II (BD). 2×10^5^ were intravenously transferred into *Rag1*^−/−^ mice that had received 4Gy of irradiation.

### Tumor cell adoptive transfer

For adoptive transfer of tumor cells, the inguinal, brachial, and axillary lymph nodes were prepared and stained as above. Tumor cells (7AAD^−^CD19^+^B220^−^IgM^−^CD93^+^) were either FACS sorted on a FACS ARIA-II (BD) or were calculated as a fraction of total cells by flow cytometry. 1×10^5^ tumor cells, either as a pure population or as a total cell suspension, were intravenously transferred into wildtype C57BL/6 mice.

### Single Cell RNA sequencing

Live pro-B cells (CD19^+^B220^+^IgM^−^IgD^−^CD24^low^CD43^+^) from *Pax5*^*wt/wt*^ and *Pax5*^*Y351*/Y351\**^, and live CD19^+^B220^−^ cells (CD19^+^B220^−^IgM^−^IgD^−^) from *Pax5*^*Y351*/Y351\**^ mice were isolated from the bone marrow by FACS sorting 50 cells into 2μl of RNA lysis buffer with 0.5μl of oligo-dT primer (5mM) and 0.5μl of dNTP mix (10mM) into 96 well V bottom plates with 4 replicates sorted per sample resulting in a total of 200 sequenced cells per sample. Plates were spun briefly to collect contents at the bottom of the well and subsequently wrapped in foil and stored at −80°C. RNA sequencing was performed using a modified version of the SMARTseq2 protocol (*39*) as described in (*40*). Sequencing was performed on the Illumina NextSeq sequencing platform.

The paired-end (PE) reads from each sample were first trimmed using the PE mode of Trimmomatic v0.36 (*41*). The trimmed reads were aligned to the mouse reference genome (NCBIM37) using STAR 2.7.0e (*42*). FeatureCounts v1.6.1 (*43*) was then applied to extract the number of mapped reads to each gene to produce the gene expression profiles. After that, DESeq2 (*44*) from the Bioconductor R package was used to perform number of differential gene expression studies.

### Whole exome sequencing

2×10^6^ tumor cells (7AAD^−^CD19^+^B220^−^IgM^−^CD93^+^) per sample were FACS sorted on a FACS ARIA I or II and whole exome sequencing and variant calling performed as previously described (*37*).

### Survival curve

GraphPad Prism 8 was used to calculate survival curves based on B-ALL development. Mice that were confirmed to have died from tumor development or if the cause of death was unknown and therefore tumor development could not be ruled out, were classified as having “died” and given a score of 1 at the time of their death. Mice that died due to unrelated causes (e.g. fight wounds, malocclusion) or were used for experiments prior to tumor development, were classified as “censored” and given a score of 0 at the time of their death. Any mouse that was still alive at the time of data collection was classified as censored.

### Gene expression comparison between wild-type and malignant cells

All samples were obtained from various sources through controlled or public access. The series from Lund University (LUND) (EGAS00001001795) (*45*) was downloaded from the European Genome-phenome Archive (EGA). The series from Children’s Hospital of Philadelphia (CHOP) (GSE115656) was downloaded from Gene Expression Omnibus (GEO) (*46*). Samples from TARGET (Therapeutically Applicable Research to Generate Effective Treatments) samples were downloaded from the TARGET data portal at National Cancer institute (NIH) together with the associated clinical information, corresponding to dbGAP accession phs000463 (ALL phase 1) and phs000464 (ALL phase2). Data from patients from the Princess Maxima Center for Pediatric Oncology (PMJCI) from (*47*) was obtained from the authors. Samples form mouse from Immgen Project (*48*) were downloaded from Immgen data portal, corresponding to GEO accession GSE109125. All the information related with the clinical information and sample extraction are described in detail in their respective publication (Supplementary Table 4).

Samples and gene expression values from human samples were mapped to the GRCh38.p10 reference human genome assembly (Gencode v27) using SALMON (v 0.7.2) at transcript level by transcripts per million (TPM). Quantification at gene level was performed using pseudo counts from SALMON quantification and transforming to counts by gene using tximport library function from Bioconductor (*49*). We used STAR-Fusion (*50*) to identify candidate chromosomal rearrangements giving rise to gene-fusions from RNA-seq data. The index was generated using the same used before. STAR-Fusion was run for each FASTQ file using the default parameters described at https://github.com/STAR-Fusion/STAR-Fusion/wiki/Home/. We selected a total of 11882 orthologous genes between human and mouse using Biomart database selecting one two one orthologous genes.

Normalization of the gene counts was perform using TMM normalization model using Limma package (*51*) in R to obtain logCPM values by gene. We removed the batch effect associated with different types of cohort using removeBatchEffect function from Limma with a design matrix related to condition tumor or normal sample.

## Acknowledgements

This study used National Collaborative Research Infrastructure Strategy -enabled Australian Phenomics Network and Bioplatforms Australia infrastructure and we thank the Australian Phenomics Facility Next Generation Sequencing team and the Biomolecular Resource Facility at the John Curtin School of Medical Research for performing whole exome and RNA sequencing. We thank the Genome Informatics group for exome sequencing analysis and the staff at the Australian Phenomics Facility for animal husbandry. We also thank the National Computational Infrastructure (Australia) for continued access to significant computation resources and technical expertise.

## Author contributions

C.M.R., A.E., and C.C.G. identified the ENU mouse strain. A.E. and B.B. designed the research. B.B., J.R., K.H., H.J.S., H.B., M.Y., N.B., S.A.O., C.M.R., C.Y., L.A.M. performed experiments. B.B., J.R., K.H., C.M.R., analyzed data. T.D.A., X.L., V.C., A.C. and E.E. analyzed RNA and WES data. S.L.N., K.M.H., and I.A.C. provided additional oversight and strategic direction. B.B., L.A.M., and A.E. wrote and edited the manuscript.

## Data and materials availability

The *Pax5*^*Y351\**^ mice are available through the Australian Phenome Bank (https://pb.apf.edu.au).

## Supplementary Materials

**Fig. S1.** PAX5^Y351*^ mutation causes slight growth delays in homozygous mice. Weight (g) of female and male mice at 25 and 50 days of age. Bars show average per group ± SEM with individual mice represented by circles. Asterisks indicate significance: ns P≥0.05, *P<0.05, **P<0.01, ***P<0.001, ****P<0.001 by two-way ANOVA with Tukey correction for multiple comparisons.

**Fig. S2.** *Pax5*^*Y351*/Y351\**^ B cells downregulate IgD but not IgM. Quantitative expression (gMFI) of IgM and IgD on the surface of FoB cells (B220^+^CD23^+^CD21/35^low^) in the spleen as determined by flow cytometry. Bars show average per group ± SEM with individual mice indicated by circles. Asterisks indicate significance: ns P≥0.05, *P<0.05, **P<0.01, ***P<0.001, ****P<0.001 by one-way ANOVA with Tukey correction for multiple comparisons. Data representative of at least three independent experiments.

**Fig. S3.** Altered B1a and B1b ratios in the peritoneal cavity of *Pax5*^*Y351*/Y351\**^ mice. **(A)** Representative FACS plots of the gating strategy for the peritoneal cavity. B1: B220^low^CD19^hi^, B1a: B220^low^CD19^hi^CD5^+^, B1b: B220^low^CD19^hi^CD5^−^. **(B)** Frequency of B cell subsets in the peritoneal cavity. Bars show average per group ± SEM with individual mice indicated by circles. Asterisks indicate significance: ns P≥0.05, *P<0.05, **P<0.01, ***P<0.001, ****P<0.001 by one-way ANOVA with Tukey correction for multiple comparisons. Data representative of at least three independent experiments.

**Fig. S4.** Mutant CD19^+^B220^−^ cells maintain high expression of PAX5 and are dissimilar from wildtype B1 and MZB cells. **(A)** Expression of PAX5 in B cell subsets in the bone marrow, as determined by flow cytometry. **(B)** Expression of CD43 on wildtype B1 cells and *Pax5*^*Y351*/Y351\**^ CD19^+^B220^−^ cells in the spleen. **(C)** Expression of CD1d and CD9 on wildtype MZB and *Pax5*^*Y351*/Y351\**^ CD19^+^B220^−^ cells in the spleen. **(A - C)** Bars show average per group ± SEM with individual mice indicated by circles. Asterisks indicate significance: ns P≥0.05, *P<0.05, **P<0.01, ***P<0.001, ****P<0.001 by one-way ANOVA with Tukey correction for multiple comparisons. Data representative of at least three independent experiments.

**Fig. S5.** PAX5^Y351*^ mutation has no effect on lymphoid and myeloid progenitor cell development. **(A)** Representative FACS plots of lymphoid and myeloid progenitor cell gating strategy in the bone marrow. **(B)** Total cell number (x10^6^) of hematopoietic stem cells (HSC), multipotent progenitors (MPP), common lymphoid progenitors (CLP), and B cell primed progenitor (BLP) subsets in the bone marrow. Bars show average per group ± SEM with individual mice indicated by circles. Asterisks indicate significance: ns P≥0.05, *P<0.05, **P<0.01, ***P<0.001, ****P<0.001 by one-way ANOVA with Tukey correction for multiple comparisons. Lin: lineage markers (CD4, CD8, TCRβ, TCRγδ, CD11b, CD11c, Ter119, NK1.1, IgD, IgM, Gr1). Data representative of at least three independent experiments.

**Fig. S6.** *Pax5*^*Y351*/Y351\**^ cells have significantly different expression of PAX5 targeted gene sets compared to wildtype but normal expression of housekeeping gene sets. Comparison of fitted reads of housekeeping and PAX5-target gene sets in wildtype pro-B cells and homozygous Pro-B and CD19^+^B220^−^ cells. Box plots show the average distribution of fitted reads, normalized to the maximum expression level per gene. Housekeeping gene sets including metabolism, mitosis, and mitochondrial obtained from the Broad Institute Gene Set List, further details listed Supplementary Table 2. n=3 per group, asterisks indicate significance: ns P≥0.05, ***P<0.001, ****P<0.001 by one-way ANOVA with Tukey correction for multiple comparisons.

**Fig. S7.** Female *Pax5*^*Y351*/Y351\**^ mice have a shorter life-expectancy compared to males. Kaplan-Meier survival curve showing death by confirmed tumor development in *Pax5*^*Y351*/Y351\**^ male (brown) and *Pax5*^*Y351*/Y351\**^ female mice (purple). Grey box shows difference between median age of survival in female (197 days) and male (247 days) mice. Asterisks indicate significance: ****P<0.001 by Log-rank (Mantel-Cox) test.

**Fig. S8.** *Pax5*^*Y351*/Y351\**^ tumors do not rearrange the immunoglobulin heavy chain. Intracellular IgM expression (cμIgH) in tumor cells obtained from *Pax5*^*Y351*/Y351\**^ mice. Histogram shows representative plot of intracellular IgM expression in wildtype precursor B cells (B220^+^IgM^−^IgD^−^) and homozygous tumor cells (CD19^+^B220^−^CD93^+^). Bar graph shows frequency of cells expressing intracellular IgM (IgH). Bars show average per group ± SEM with individual mice indicated by circles. Asterisks indicate significance: ***P<0.001 by students two-tailed T test. Data representative of at least two independent experiments.

**Table S1.** RNAseq of PAX5 Target Genes

**Table S2.** RNAseq of Housekeeping Gene sets

**Table S3.** Source information for data used to compare RNAseq of wildtype and tumor cells

**Table S4.** Whole Exome Sequencing of *Y351*/Y351\** tumor cells.

