## Supplementary figures and images for "Dysregulation of PAX5 causes uncommitted B cell development and tumorigenesis in mice"

### Fig. S1

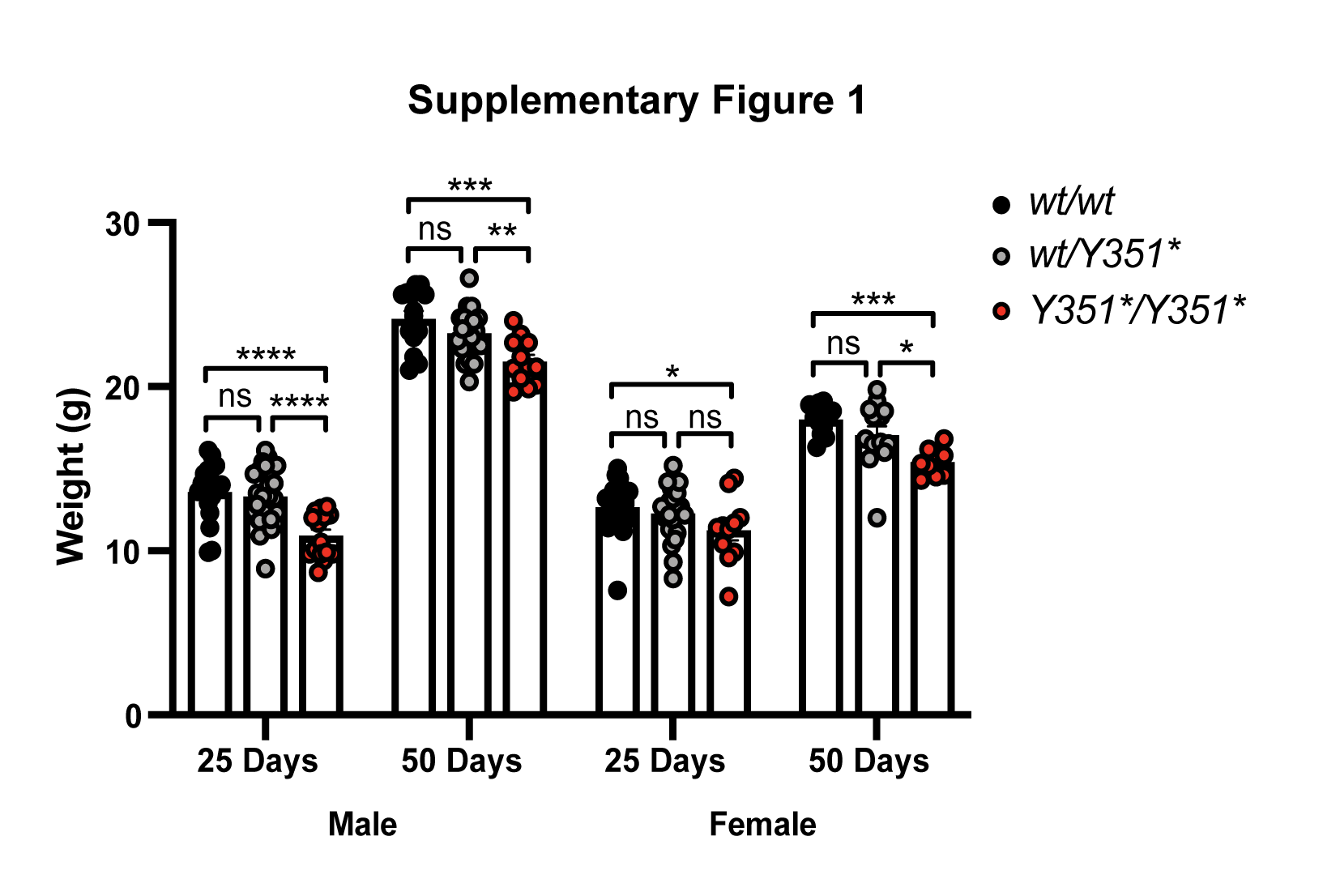

### Fig. S2

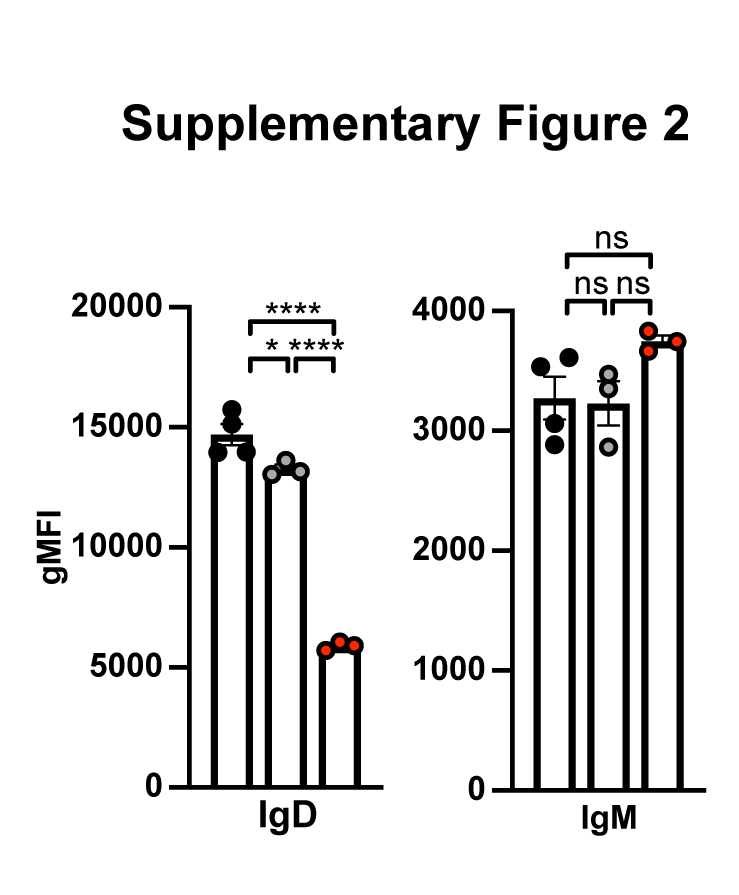

### Fig. S3

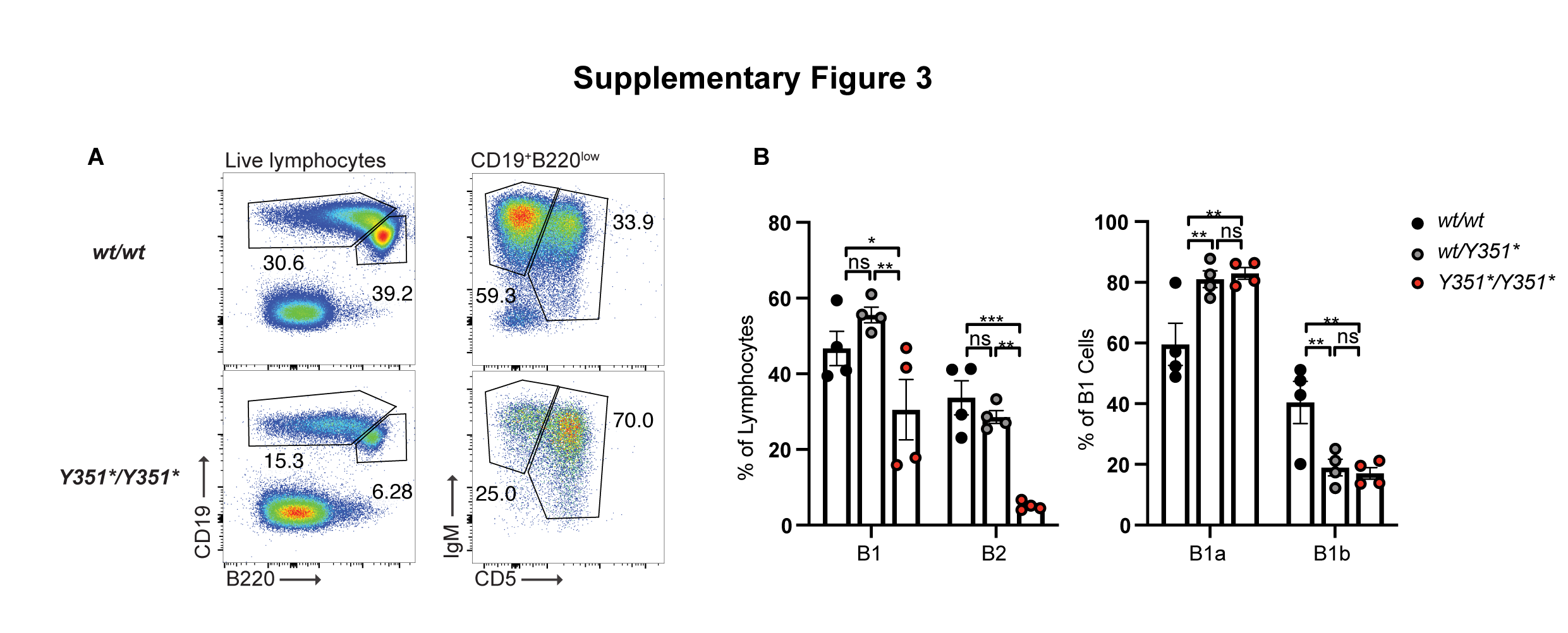

### Fig. S4

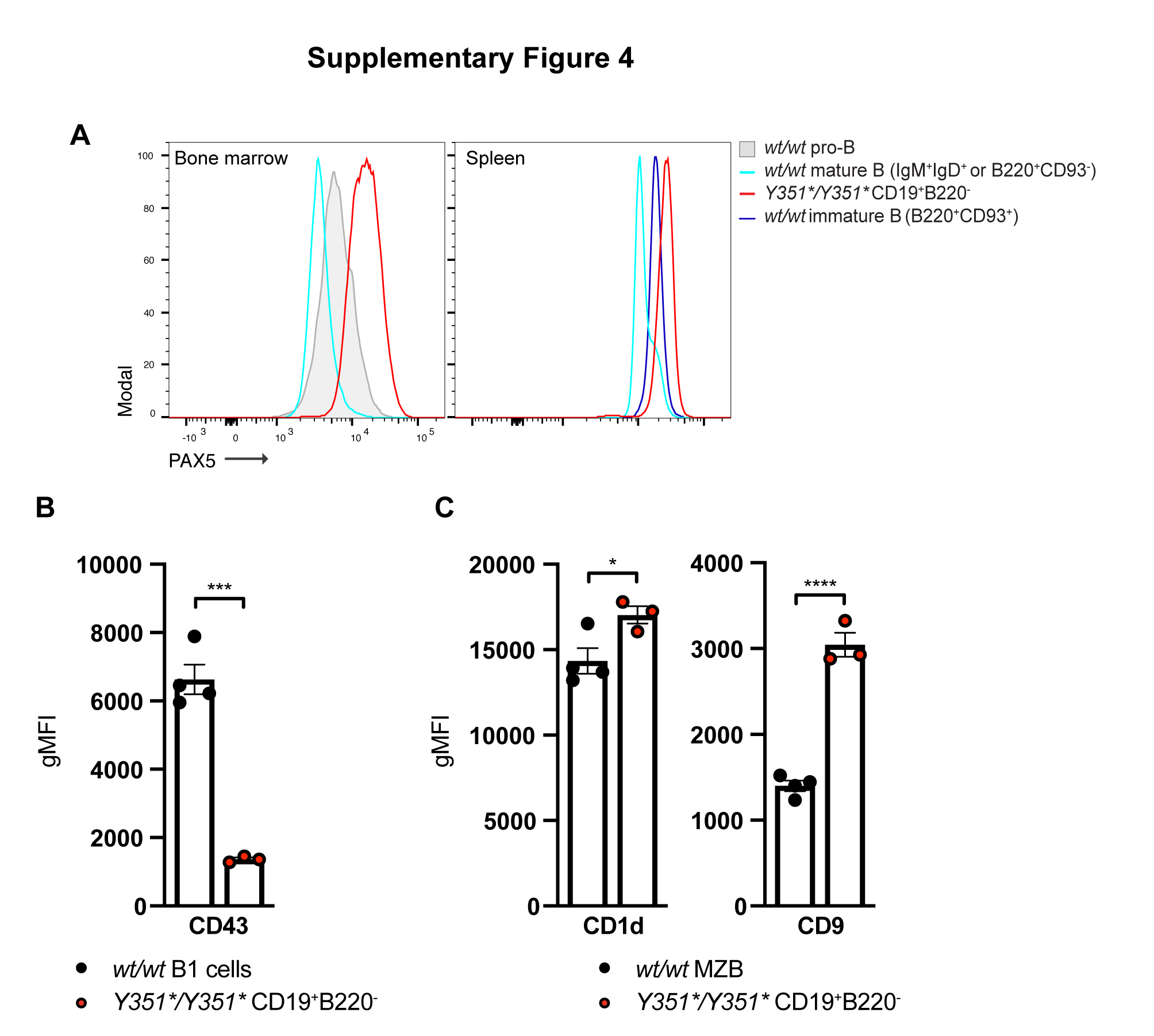

### Fig. S5

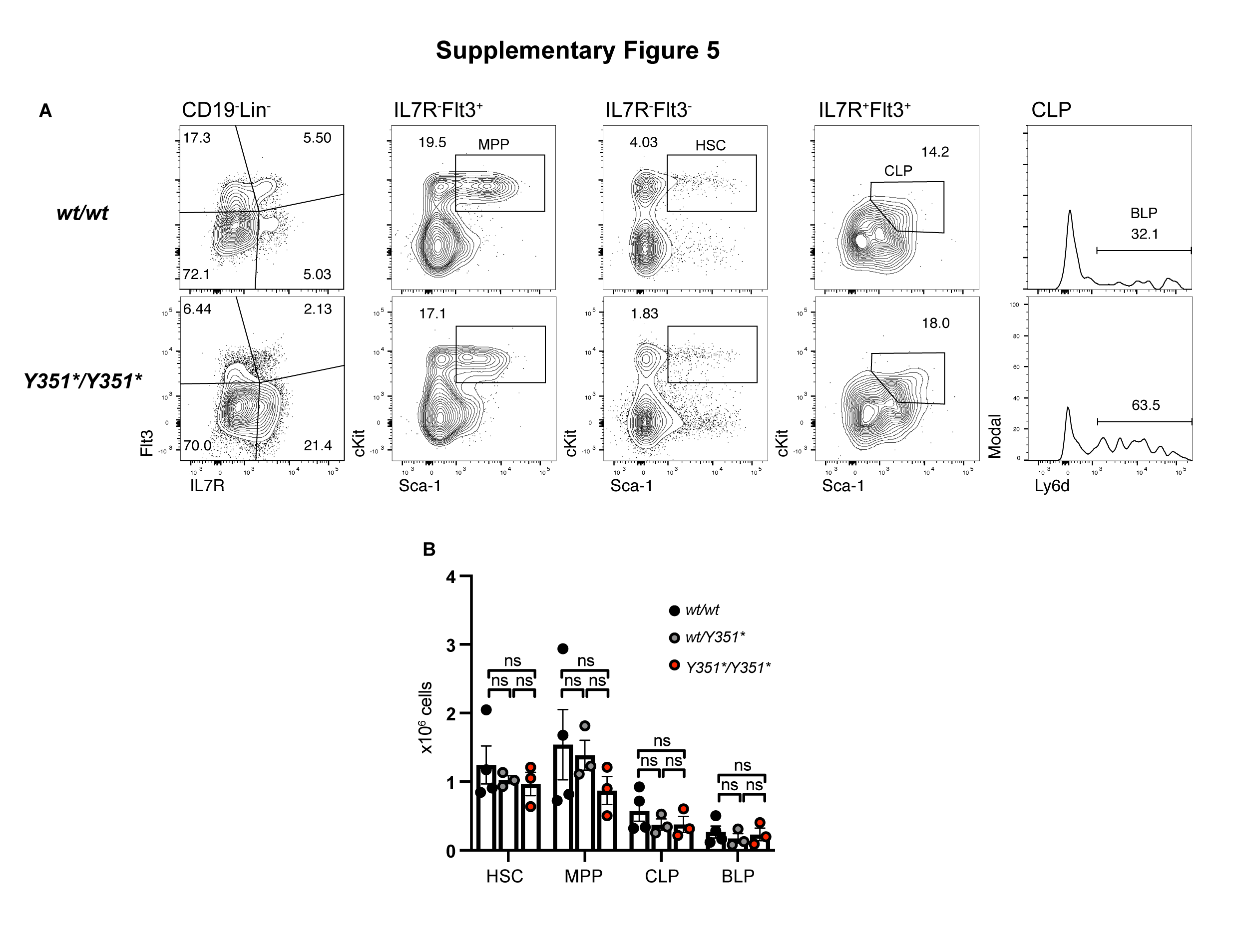

### Fig. S6

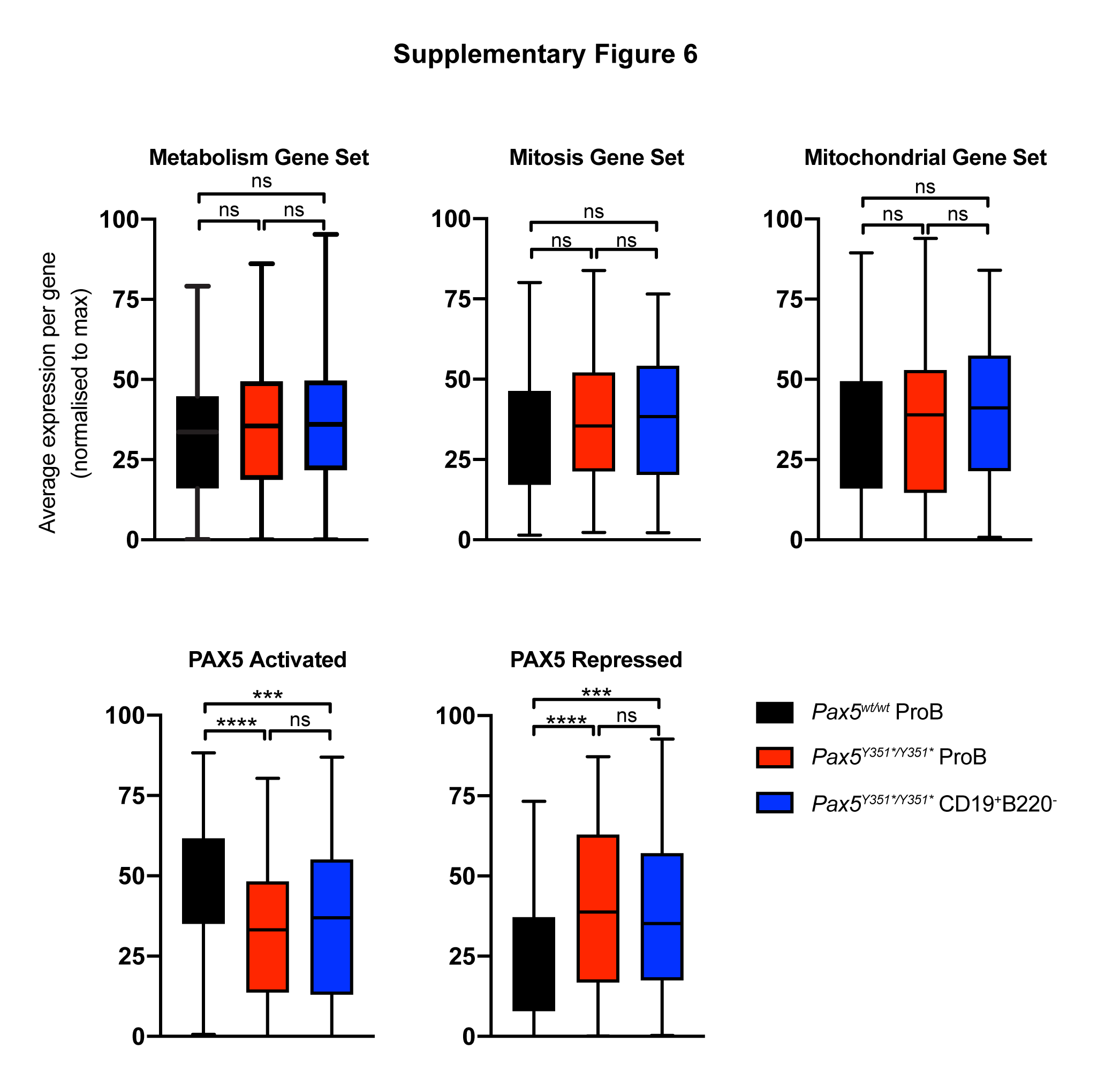

### Fig. S7

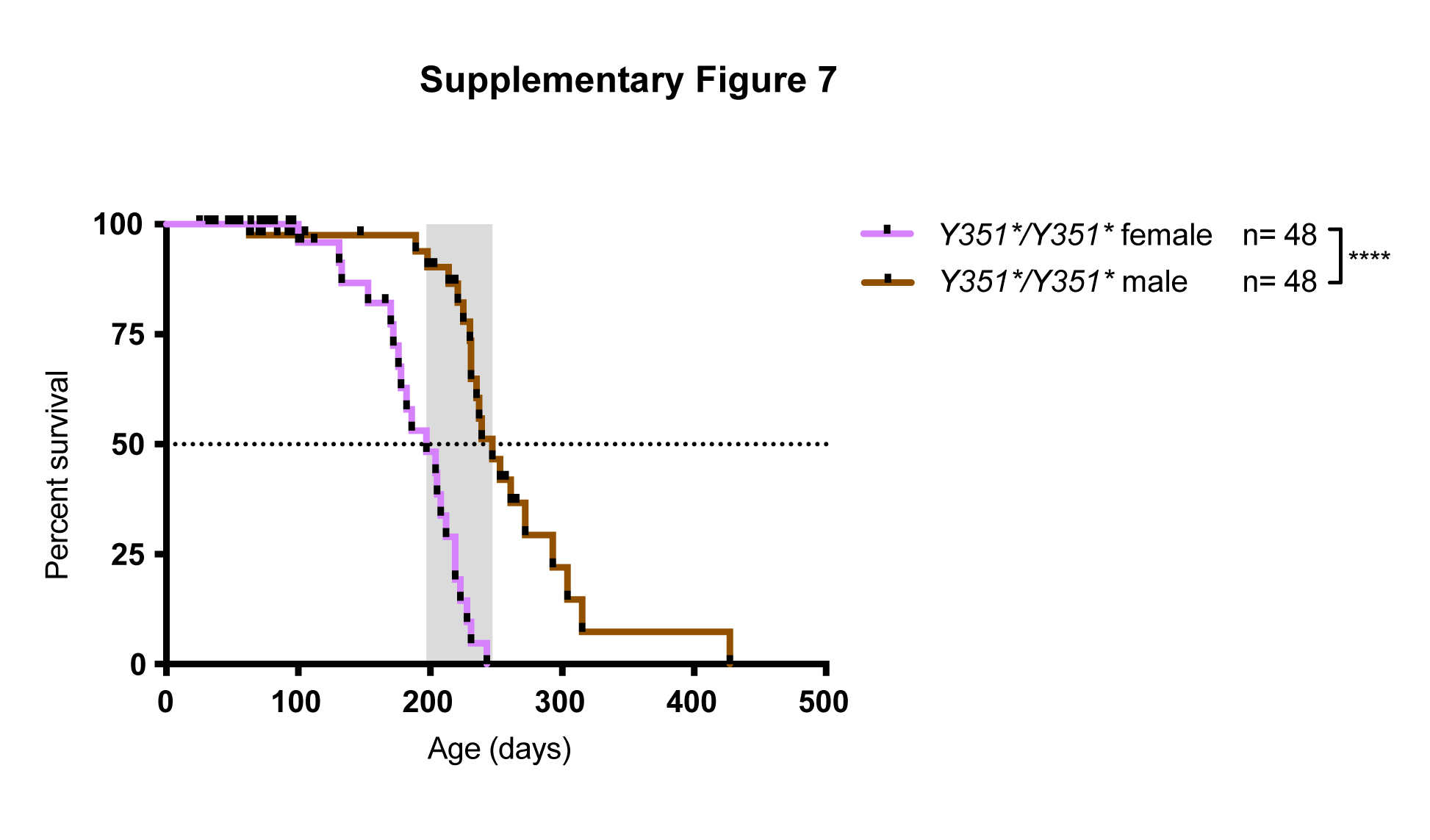

### Fig. S8

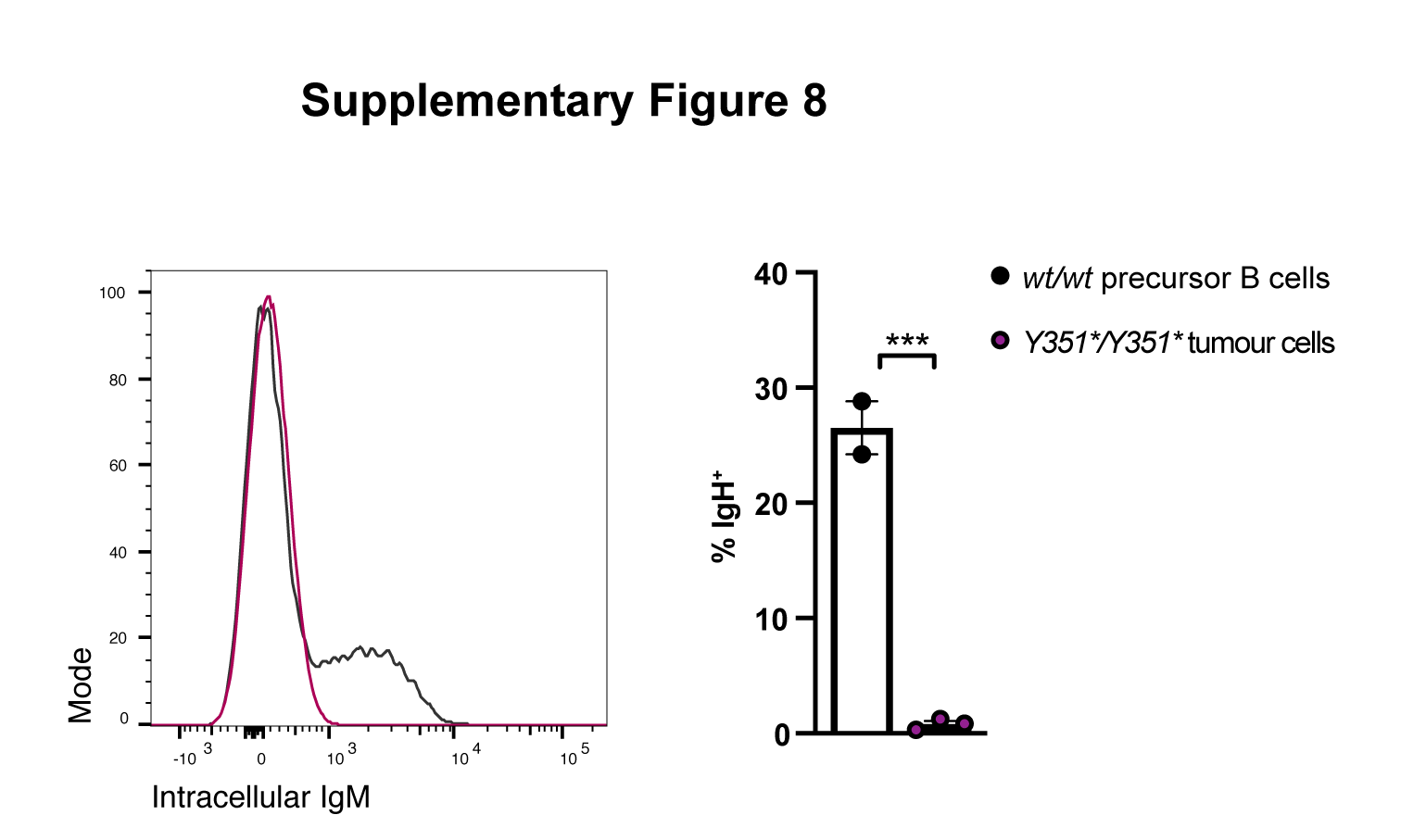
